# Cardiorespiratory and Cardiac Biomarker Responses to Five Anesthetic Regimens in Rats

**DOI:** 10.64898/2026.04.07.716572

**Authors:** Larissa de Jesus Corrêa, Vítor Sampaio Minassa, Bianca Teixeira Jara, Bruno Antonio Alday De Moura, Thatiany Jardim Batista, Juliana Barbosa Coitinho, Daniela Amorim Melgaço Guimarães do Bem, Leonardo dos Santos, Julian Francis Richmond Paton, Fiona Dianne McBryde, Vanessa Beijamini, Igor Simões Assunção Felippe, Karla Nívea Sampaio

**Affiliations:** Postgraduate Program in Pharmaceutical Sciences, Department of Pharmaceutical Sciences, Federal University of Espírito Santo, Vitória, ES, Brazil; Postgraduate Program in Physiological Sciences, Department of Physiology, Federal University of Espírito Santo, Vitória, ES, Brazil; Postgraduate Program in Biochemistry, Department of Physiology, Federal University of Espírito Santo, Vitória, ES, Brazil; Department of Pharmaceutical Sciences, Federal University of Espírito Santo, Vitória, ES, Brazil; The Centre for Heart Research – Manaaki Mānawa, Department of Physiology, Faculty of Health & Medical Sciences, University of Auckland, Grafton Campus, Auckland, 1023, New Zealand

**Author notes:** Corresponding authors: **Professor Karla Nívea Sampaio**: Federal University of Espírito Santo; Postgraduate Program in Pharmaceutical Sciences; Department of Pharmaceutical Sciences, Health Sciences Center; Av. Marechal Campos, 1468, Maruípe, Vitória, ES, Brazil; Postcode: 29047-105; **Dr. Igor Simões Assunção Felippe**: The Centre for Heart Research – Manaaki Mānawa, Department of Physiology, Faculty of Health & Medical Sciences, University of Auckland, Grafton Campus, Auckland, 1023, New Zealand. L.J.C., V.S.M and B.T.J. are equal first authors.

**Keywords:** general anesthesia, cardiorespiratory, biochemical, cardiac biomarkers

## Abstract

General anesthetics enable invasive experimentation but can affect cardiovascular and respiratory physiology, biasing preclinical outcomes. We compared five anesthetic regimens in adult male Wistar rats, tribromoethanol (TBE, 250 mg/kg i.p.), chloral hydrate (CH, 400 mg/kg i.p.), ketamine–xylazine (KX, 80/10 mg/kg i.p.), thiopental (TP, 80 mg/kg i.p.), and isoflurane (ISO, 4% induction, 2% maintenance), to investigate integrated cardiorespiratory and biochemical markers. Femoral arterial catheterization allowed continuous blood pressure (BP) and derived heart rate (HR) recordings, while ventilation was assessed through pletysmography at baseline (awake), during induction, and recovery phases of anesthesia. Variability was evaluated in the time and frequency domains, including HR, systolic blood pressure (SBP), and spontaneous baroreflex sensitivity. In an independent cohort of rats, butyrylcholinesterase (BChE), CK-MB, cTnI, and LDH were measured. Baseline BP was unchanged by TBE and TP, whereas all anesthetics affected HR. Minute ventilation and breathing frequency were reduced with all agents, while tidal volume decreased with KX and TBE only. LDH and cTnI were unaffected, BChE was reduced by KX, TBE, and ISO, and CK-MB increased with CH and KX. Variability analysis showed that all anesthetics depressed pulse-interval and SBP variability and shifted spectral power toward higher frequencies, while baroreflex sensitivity and effectiveness were consistently reduced. During recovery, KX and TP restored most variability indices, whereas CH, TBE, and ISO showed persistent suppression. These findings highlight distinct profiles of cardiovascular depression and biomarker responses across anesthetics and underscore the importance of accounting for autonomic variability when selecting different anesthetics in experimental protocols.

**Highlights:**

- Five anesthetic regimens were tested in rats.
- All anesthetics reduced ventilation, and KX and TBE also reduced tidal volume.
- CH and KX increased CKMB, while KX, TBE and ISO reduced BChE.
- All anesthetics reduced blood pressure variability and baroreflex sensitivity.
- Variability recovered with TP and KX, whereas CH, TBE and ISO showed persistent suppression.

## 1. Introduction

General anesthesia (GA) is a critical intervention in ensuring patients remain insensitive to pain during invasive procedures in both clinical and preclinical settings (P. Flecknell, 2016; Franks, 2008). An ideal anesthetic agent should rapidly and safely induce unconsciousness (i.e., hypnosis), provide effective analgesia, cause minimal side effects, and allow for quick recovery from amnesia (Brown et al., 2011; P. A. Flecknell, 2009a).

Although the underlying mechanism of GA is not completely understood, a general conceptual framework suggests that its hypnotic and analgesic effects stem from an overall depression of the central nervous system (CNS), achieved via both a reduction in excitatory synaptic transmission and an increase in inhibitory signaling (Franks, 2008; Hemmings et al., 2019). The main pharmacological classes of general anesthetic agents tend to modulate neurotransmission through GABA_A_, glycine, N-methyl-D-aspartate (NMDA), and nicotinic cholinergic receptors (Franks, 2008; Hemmings et al., 2019). However, given the ubiquitous nature of the aforementioned drug targets, their modulation can significantly affect other physiological systems, including the cardiovascular and respiratory systems (Albrecht et al., 2014; Eger, 2004; Saraswat, 2015).

There is evidence in the literature indicating that anesthetic agents can persistently interfere with cardiorespiratory parameters in both humans and animals (Albrecht et al., 2014; Eger, 2004; P. A. Flecknell, 2009b; Heistein, 2006; Khan et al., 2014; Saraswat, 2015). For instance, Albrecht et al. (2014) investigated the effects of repeated anesthesia using different agents and reported that isoflurane (ISO) induces reductions in blood pressure (BP) and heart rate (HR) in Wistar rats. Notably, the depressant effect became more pronounced with an increased number of anesthetic exposures (Albrecht et al., 2014) Likewise, a combination of medetomidine-midazolam-fentanyl significantly decreased animals’ pulse pressure whilst ketamine-xylazine (KX) was associated with reductions in sleeping time and body weight (Albrecht et al., 2014).

The use of animal models in preclinical studies is essential for advancing scientific knowledge in research fields such as cardiovascular and respiratory physiology (Mosneag et al., 2024; Olmos-Pastoresa et al., 2023; Parikh & Pierce, 2024; Shrestha et al., 2023; van Doorn et al., 2024). Consequently, the administration of anesthetic agents is common practice in experimental settings due to the invasive nature of many procedures (Gargiulo et al., 2025; Maud et al., 2014; Navarro et al., 2021; Oh & Narver, 2024; Pachon et al., 2015). The choice of the anesthetic agent, including the route of administration, is typically guided by the species involved and the specific aims of the study (P. Flecknell, 2016; Gargiulo et al., 2012, 2025; Oh & Narver, 2024). However, despite the critical role anesthetics play in hypothesis-driven research, little is known about their potential to alter commonly measured physiological parameters, such as blood biomarkers, in the days following the procedure (Deckardt et al., 2007; Ochiai et al., 2016). Therefore, the choice of anesthesia may represent an important confounding variable, potentially contributing to discrepancies observed in the literature (P. Flecknell, 2016; Morton et al., 2019; Mosneag et al., 2024).

In cardiovascular studies, cardiac troponin I (cTnI), creatine kinase-MB fraction (CK-MB), lactate dehydrogenase (LDH) and butyrylcholinesterase (BChE) are commonly used to diagnose cardiac injury. cTnI is a highly specific marker for cardiac tissue, allowing the detection of even small injuries for up to 4 hours after myocardial damage (Danese & Montagnana, 2016; Garg et al., 2017; Létienne et al., 2006). CK-MB is a less specific marker of cardiac injury. Although it is more concentrated in the cardiac muscle, it is also found in skeletal muscle, making it a useful but not exclusive indicator of myocardial infarction. Similarly, LDH is an enzyme found in various tissues and is released during tissue degradation (Danese & Montagnana, 2016). Interestingly, recent studies have also demonstrated that BChE activity is an important enzyme to monitor, as it holds prognostic value for mortality in patients with acute myocardial infarction (Brzezinski-Sinai et al., 2021; Michels et al., 2021; Sun et al., 2016).

Herein we systematically evaluated the effects of five commonly used anesthetic agents, i.e., ISO, Chloral Hydrate (CH), KX, Thiopental (TP), and Tribromoethanol (TBE), on cardiorespiratory parameters and cardiac injury biomarkers. By integrating functional and molecular assessments, this study aimed to characterize the physiological impact of anesthesia, offering valuable insights to guide anesthetic selection in preclinical research and helping to minimize variability in physiological outcomes across experimental studies.

## 2. Methods

### 2.1. Ethical approval

All experiments complied with the Guide for the Care and Use of Laboratory Animals published by the Brazilian National Council for Animal Experimentation Control, with the ARRIVE guidelines, and with the Directive 2010/63/EU of the European Parliament on the protection of animals used for scientific purposes. The Institutional Ethics Committee approved all experimental protocols for animal experimentation at the Federal University of Espírito Santo (CEUA-UFES; Approval protocols numbers: 29/2015 and 17/2023). All steps to minimise the animals’ pain and suffering have been taken. The investigators understand the ethical principles under which the journal operates, and the work presented here complies with it.

### 2.2. Animals and Housing

A total of (n=100) male Wistar rats (300-350 g) were obtained from the Animal Care Facility of the Center of Health Sciences at the Federal University of Espírito Santo, Espírito Santo, Brazil. Rats were housed in groups of 4 per cage (49 x 34 x 26 cm) in a temperature (22 ± 2 °C) and humidity-controlled room with a 12 h light/dark cycle (lights on at 6:30 a.m.). Standard chow and tap water were provided *ad libitum*. Cages were lined with wood shavings, and cleaning with bedding replacement being performed 3 times/week.

### 2.3. Anesthetic agents, dosing regimens, and preparation of solutions

Dose selection of anesthetics was carefully determined based on evidence from previous studies. Injectable anesthetics were administered intraperitoneally (i.p.) at the following doses: sodium TP (Cristália; 80 mg/kg), CH (VETEC; 400 mg/kg), KX (Synthec, Brazil; 80 mg/kg and 10 mg/kg, respectively), and TBE (Sigma-Aldrich, USA; 250 mg/kg) (Gonca, 2015; Herbst et al., 2019). The inhalant anesthetic ISO (Cristália, SP) was delivered using a vaporizer (i.e., 4% ISO for induction and 2% ISO for maintenance in O_2_ at 1L/min) as previously described (Herbst et al., 2019).

Non-commercial injectable anesthetic solutions were prepared following specific recommendations to ensure safety and reduce complications (P. Flecknell, 2016; Lieggi et al., 2005; Meyer & Fish, 2005a; Vachon et al., 2000; Weiss & Zimmermann, 1999; Zatroch et al., 2016). Tribromoethanol was dissolved in saline at 25 mg/mL (250 mg/kg at 10 mL/kg), vortexed for 5 min at 42°C, filtered, and stored in amber-protected vials at 5 °C for up to two weeks, as recommended to minimize peritonitis and chemical degradation (Lieggi et al., 2005; Meyer & Fish, 2005a). Chloral hydrate was prepared at 40 mg/mL (400 mg/kg at 10 mL/kg), vortexed for 2 min at room temperature, filtered, and stored under light protection at 5°C for up to two weeks. The use of low-concentration, high-volume solutions has been shown to lessen peritoneal irritation (Vachon et al., 2000). Because of its aqueous instability, thiopental was freshly prepared immediately before administration by dissolving crystalline powder in saline to deliver the 80 mg/kg dose, following refinement guidelines for i.p. injection (P. Flecknell, 2016; Zatroch et al., 2016).

Although the use of chloral hydrate is contraindicated by recent international guidelines due to its narrow therapeutic index and potential adverse effects, it remains in use in the preclinical setting, as evidenced by some recent studies that adopted it as an anesthetic agent (Costas-Ferreira et al., 2023; Gargiulo et al., 2025; Keshavarz et al., 2023; Shcherbak et al., 2024; Ward-Flanagan & Dickson, 2023; Zhang et al., 2023). Its inclusion in the present work was authorized by the Institutional Animal Care and Use Committee (CEUA-UFES; protocols 29/2015 and 17/2023) for methodological comparison.

### 2.4. Experimental design

Rats were randomly allocated to one of the five experimental groups, based on the anesthetic that was selected, using Excel’s RAND function. Two independent cohorts were employed: one for cardiorespiratory assessments (Protocol 1; N=60) and another for biochemical assays (Protocol 2; N=40).

The cardiorespiratory parameters were measured during both the *pre-anesthetic*, *induction*, and *recovery* phases of anesthesia (Protocol 1), which were defined based on standard reflex responses used to assess anesthetic depth and monitoring periods (Albrecht et al., 2014). On the first day, rats underwent catheterization surgery for cannulation of the femoral artery and vein. Anesthesia was administered using the assigned agent, as described above. On the second day, 24 hours after recovery, baseline cardiorespiratory parameters were recorded for 30 minutes in freely moving conscious rats; this period was defined as the *pre-anesthetic* phase. After this period, rats were re-anesthetized, and the *induction* phase was defined as the first 10 minutes after loss of pedal reflex, during which both BP and HR were continuously monitored. Monitoring was maintained until the return of righting reflex, which defines the end of anesthesia. Then, *recovery* phase was defined as the final 10 minutes preceding the return of righting reflex. For anesthesia using ISO, after induction, animals were maintained sedated for 40 minutes before the vaporizer was turned off, in order to simulate the duration of sedation evoked by injectable agents.

The effects of anesthesia on cardiac injury biomarkers were evaluated only during the *pre-anesthetic* and *induction* phases (Protocol 2). For this protocol, rats underwent the same cannulation surgery with 24 hours recovery period. Blood sampling (0.7 mL) was performed in conscious rats via the arterial line (Figure 1).

**Figure 1.**
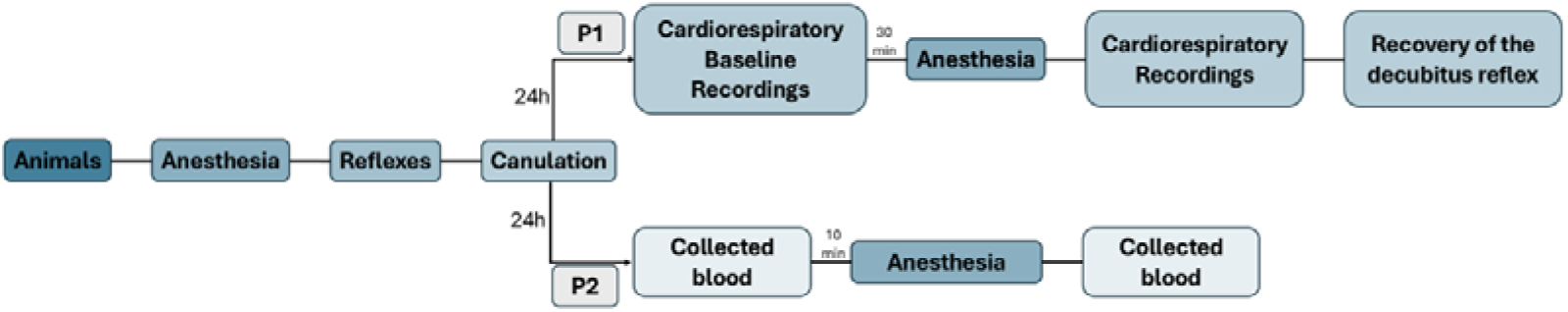
Experimental timeline showing the protocols performed for cardiorespiratory and biochemical assays.

### 2.5. Catheterization surgery

Under anesthesia, rats were given a single subcutaneous injection of analgesic (metamizole - 250 mg/kg). In addition, animals were kept warmed (∼ 37°C) by wrapping the animal in cotton wool or by using the bubble packing to prevent heat loss (Flecknell, 2009b), petroleum-based artificial tear ointment was applied to the eyes to maintain corneal lubrication, and warmed saline (0.9% NaCl, 5 mL/kg, 35–37°C) was administered subcutaneously to ensure hydration.

Surgical fields were trimmed and disinfected with iodinated alcohol solution. Under aseptic conditions, the right femoral vein and artery were exposed via a 1 cm incision in the inguinal region. Each vascular line consisted of polyethylene catheters (PE-10 connected to PE-20), pre-filled with heparinized saline (50 U/mL). Catheters were inserted approximately 1.5 cm into the vessels, tunneled subcutaneously, and exteriorized through the back of the neck.

Following surgery, animals were warmed under a heating lamp for 10 minutes. Postoperative analgesia (metamizole, 250 mg/kg) was administered subcutaneously every six hours. Animals were then allowed to recover for 24 hours in individual cages with *ad libitum* access to food and water.

### 2.6. Assessment of anesthesia induction

During anesthesia, onset of hypnosis was confirmed by assessing the palpebral and righting reflexes, as well as maintenance of dorsal recumbency, whereas depth of anesthesia was assessed via tail pinch and pedal reflex. For ISO, only the righting reflex was assessed at this phase, as induction occurred in a sealed chamber, precluding testing other reflexes. The time for reflexes loss and duration of anesthesia was recorded.

### 2.7. Cardiorespiratory Parameters

For BP recordings, the arterial line was connected to a pressure transducer (MLT0670, ADInstruments, Australia), while signals were amplified (BP Amp, ADInstruments, Australia), digitized (Powerlab, ADInstruments, Australia), and saved using LabChart8 software (ADInstruments, Australia). In freely moving conscious rats, systolic (SBP), diastolic (DBP) and mean arterial pressures (MAP, in mmHg) were recorded while HR (in beats-per-minutes) and pulse interval (PI, in milliseconds) were derived from pulsatile arterial pressure (PAP).

For respiratory recordings, the barometric method of Drorbaugh & Fenn (Drorbaugh & Fenn, 1955) was used to calculate tidal volume (VT), in mg/kg using the whole-body plethysmography. The respiratory frequency (*f_R_*), expressed as number of cycles per minute (cpm), was derived from pressure oscillations in the chamber, whilst minute ventilation (VE), in mg/kg/min, was calculated as the product of *f_R_* and V_T_, as previously described (Felippe et al., 2020).

### 2.8. Pre-anesthetic Phase

For breathing recordings, the plethysmograph chamber was sealed for no longer than two minutes per session. A minimum recovery period of 10 minutes was allowed between each recording. PAP, MAP, and HR were recorded continuously for 30 minutes in conscious, freely moving animals at rest. Data were averaged from a 10-minute segment of stable, noise-free BP, designated as the control period. ISO-treated rats did not undergo respiratory recordings under anesthesia, as mask delivery precluded accurate chamber measurements.

### 2.9. Induction Phase

Following control recordings of cardiorespiratory parameters, anesthesia was administered according to group assignment. Changes in BP and breathing activity were monitored as previously described. Ten minutes after *induction,* anesthetic depth was assessed every five minutes using a tail pinch (5-10 seconds) applied with a clip at the distal end of the tail. Cardiorespiratory parameters were continuously monitored until the return of the righting reflex.

For ISO, *induction* was achieved by saturating the plethysmograph chamber with 4% ISO in O_2_ (1L/min). After three minutes, animals were transferred to a nose cone and maintained at 2% ISO in O_2_ (1L/min). Anesthetic depth was assessed via tail pinch and visual inspection of breathing patterns. As previously mentioned, ISO was maintained for 40 minutes before the vaporizer was turned off, and the righting reflex was used to confirm recovery.

### 2.10. Cardiovascular variability

SBP and HR variability (HRV) were analyzed offline from BP recordings using CardioSeries software (version 2.3 -http://www.danielpenteado.com) as previously described (Batista et al., 2019; Felippe et al., 2020). In the time-domain, total variance was calculated for PI and SBP, along with the root mean square of successive differences (RMSSD) of inter-beat intervals.

HRV in the frequency-domain was assessed via spectral analysis. Beat-to-beat series were interpolated at 10 Hz and segmented into half-overlapping windows containing 512 data points. Nonstationary or aberrant segments, such as those with signal interruptions or movement artifacts, were visually identified and excluded from analysis. Spectral power was computed using a Fast Fourier Transform (FFT) and integrated into very low-frequency (VLF: 0–0.2 Hz), low-frequency (LF: 0.20–0.75 Hz) and high-frequency (HF: 0.75–3.00 Hz) bands. Results are presented in both absolute and normalized units. Because HF components reflect parasympathetic modulation and LF components represent a combination of sympathetic and parasympathetic influences, the LF/HF ratio was also calculated to assess sympathovagal balance.

### 2.11. Spontaneous baroreflex sensitivity

Spontaneous baroreflex sensitivity (sBRS) was evaluated using the sequence method, as previously described by Fazan et al. (2011). Baroreflex gain was calculated from upward, downward, and combined arterial pressure sequences identified in beat-to-beat data, consisting of at least four consecutive pulses in which changes in SBP were linearly correlated with changes in PI, with a correlation coefficient > 0.8. The slope of the linear regression for each sequence type was used to determine sBRS.

Because changes in SBP sequences are not always accompanied by baroreflex-mediated PI responses, the baroreflex effectiveness index (BEI) was also calculated. BEI is defined as the ratio between the number of baroreflex sequences and the total number of SBP sequences within a given time window. Both sBRS and BEI were automatically computed using CardioSeries software.

### 2.12. Blood Sampling

Samples were collected in heparinized plastic microtubes. Samples for blood analysis were kept in ice and immediately taken for analysis, whereas plasma samples for enzymatic assays were obtained by centrifugation at 1792 G for 10 min at 4 °C (SL-5 AM - Spinlab Scientific, South Korea) and stored at −80 °C.

### 2.13. Arterial blood gas analysis

Arterial blood gas analysis was performed in *pre-anesthetic* and *induction* phases using an automated blood gas analyzer at the Tommasi laboratory (GEM Premier model 3500; Instrumentation Laboratory Werfen; Bedford, MA, USA). The following parameters were measured: pH, arterial partial pressure of oxygen (PaO_2_) and carbon dioxide (PaCO_2_), bicarbonate (HCO_3_^−^), oxygen saturation (SatO_2_ in %), and total carbon dioxide (Total CO_2_).

### 2.14. Biochemistry assays

Plasma butyrylcholinesterase (BChE), creatine kinase-MB (CK-MB), cardiac troponin I (cTnI), and lactate dehydrogenase (LDH) were measured using commercially available kits (Bioclin K094-1.1, Quibasa, MG, Brazil; VITROS CK-MB, LDH, and hs Troponin I, Ortho Clinical Diagnostics, USA), following the manufacturer’s recommendations. BChE activity, expressed as U/L, was determined by a kinetic colorimetric method based on Schmidt et al., 1992. CK-MB activity, expressed as U/L, was measured by an immunoinhibition UV method according to the Scandinavian Committee on Enzymes (Hørder et al., 1979). LDH activity, in U/L, was assessed by an optimized UV kinetic method according to (Schumann et al., 2002). Cardiac troponin I, in ng/L, was determined by a high-sensitivity chemiluminescent immunoassay, following IFCC recommendations for cardiac biomarkers (Thygesen et al., 2007).

### 2.15. Statistical Analysis

Graphic and statistical analyses were performed using GraphPad Prism (version 8.0.2 for Windows; GraphPad Software®, San Diego, California, USA). Data were assessed for normality using standard normality tests and were considered normally distributed when at least one test indicated normality. The dependent variables, time to loss of reflexes during induction and time to recovery from anesthesia, were analyzed using one-way ANOVA. Cardiorespiratory parameters were assessed using one-way repeated measures ANOVA. For repeated measures ANOVA, sphericity was assessed and, when violated, Greenhouse–Geisser correction was applied. Enzyme activity/concentration and blood gas analyses were evaluated using paired t-tests and one-way ANOVA, as appropriate. When statistical significance was detected, post hoc comparisons were conducted using Tukey’s test or Dunnett’s test, depending on the experimental design. Statistical significance was set at 5%, although marginal p-values (0.05 < P < 0.08) were also reported when relevant to the interpretation of results. Data are presented as mean ± standard deviation (SD). A statistical summary table reporting exact p-values is provided in Appendix 1 at the end of the manuscript, after the References section. The final sample size (N) differs slightly across some analyses due to the exclusion of outliers identified through formal outlier tests (ROUT test, Q=1%) and animals with cannulation-related loss of cardiovascular recordings. In paired analyses, exclusions were consistently applied across all comparisons, whereas this was not required for independent analyses.

## 3. Results

### 3.1. Time to induction and recovery from anesthesia

An initial analysis of the time to loss of reflexes during *induction* (Table 1) revealed significant anesthetic-specific differences. For instance, the time to onset of hypnosis, i.e., loss of righting reflex, was significantly longer with TP compared to all other anesthetic agents (p<0.05), whereas TBE resulted in the shortest time (p<0.05). Loss of the palpebral reflex occurred sooner with TBE than TP (p<0.05), although no statistical differences were observed when compared to the other anesthetics agents. Depth of anesthesia assessed via abolition of the pedal reflex also took longer with TP than with any other agent (p<0.05). Additionally, CH was associated with a significantly longer time to loss of the pedal reflex compared to both KX and TBE (p<0.05). The loss of tail pinch reflex followed a similar pattern to that observed for the righting and pedal reflexes, with TP requiring the longest time (p<0.05), followed by CH (p<0.05).

**Table 1.**
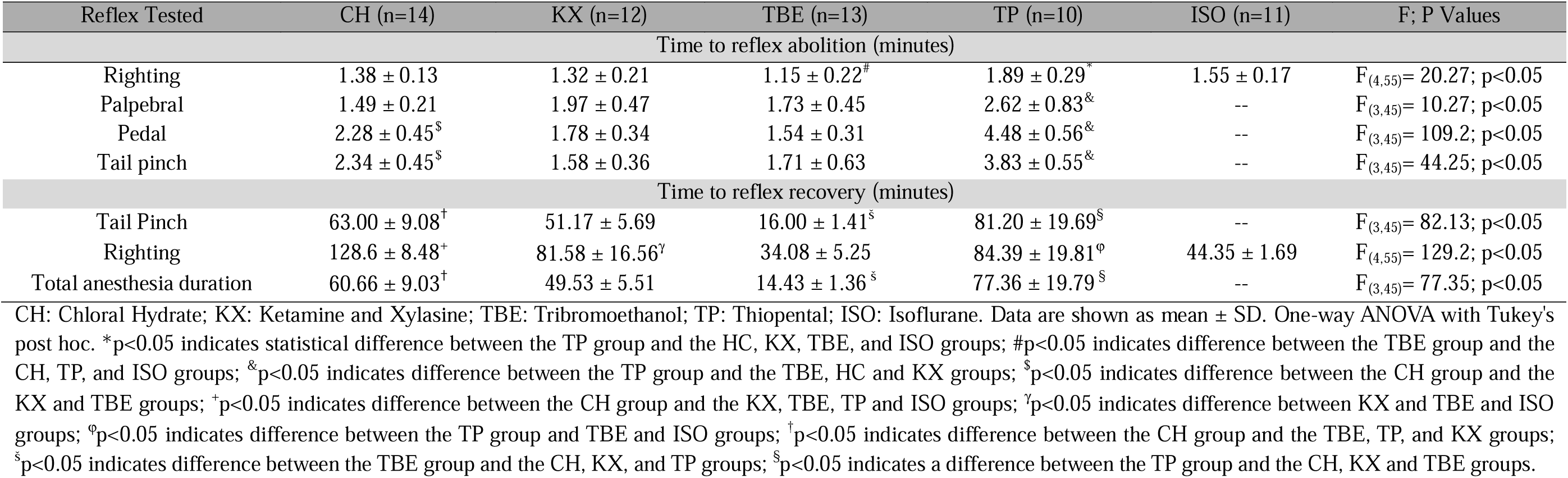
Time spent for reflexes loss and recovery, in minutes, after induction of Anesthesia with Chloral Hydrate (CH), Ketamine/Xylazine (KX), Tribromoethanol (TBE), Thiopental (TP) and Isoflurane (ISO) in rats.

As previously mentioned, the recovery from anesthesia was assessed via tail pinch and righting reflexes. TBE produced the fastest recovery of both reflexes compared to all other anesthetics agent (p<0.05).

When considering the total duration of anesthesia, defined as the interval between loss and recovery of reflexes, TBE again showed the shortest duration (p < 0.05 versus all other agents), while TP resulted in the longest (p < 0.05).

### 3.2. Cardiovascular Parameters

#### 3.2.1. Hemodynamic Data

Hemodynamic data across all phases of anesthesia are presented in Table 2. Administration of CH significantly reduced baseline SBP, DBP, and MAP ten minutes after induction (p<0.05); these parameters remained significantly lower during the *recovery* phase (p<0.05). In contrast, baseline HR increased ten minutes after induction (p<0.05), but returned to baseline levels during the *recovery* phase (p>0.05).

**Table 2.** Hemodynamic data during *pre-anesthetic*, *induction*, and *recovery* phases of anesthesia with Chloral Hydrate (CH), Ketamine/Xylazine (KX), Tribromoethanol (TBE), Thiopental (TP), and Isoflurane (ISO) in rats.

|  | CH (n=12) | KX (n=11) | TBE (n=12) | TP (n=10) | ISO (n=11) |
| --- | --- | --- | --- | --- | --- |
| SBP (mm Hg) |  |  |  |  |  |
| F; P Values | $F_{(1.968,21.65)} = 53.68$;<br>p<0.05 | $F_{(1.619,16.19)} = 20.46$;<br>p<0.05 | $F_{(1.417,15.59)} = 1.94$;<br>p>0.05 | $F_{(1.431,12.88)} = 0.65$;<br>p>0.05 | $F_{(1.735,17.35)} = 57.11$;<br>p<0.05 |
| *pre-anesthetic* | 123.6 ± 6.29 | 107.8 ± 11.92 | 115.4 ± 8.68 | 113.9 ± 10.51 | 128.1 ± 12.90 |
| *Induction* | 96.2 ± 10.52* | 152.4 ± 28.45* | 114.1 ± 14.40 | 116.8 ± 10.61 | 89.8 ± 8.88* |
| *Recovery* | 90.7 ± 10.68* | 114.0 ± 23.92 | 120.0 ± 12.21 | 117.0 ± 11.62 | 117.5 ± 9.89 |
| DBP (mm Hg) |  |  |  |  |  |
| F; P Values | $F_{(1.871,20.58)} = 76.17$;<br>p<0.05 | $F_{(1.680,16.80)} = 17.06$;<br>p<0.05 | $F_{(1.493,16.43)} = 3.40$;<br>p>0.05 | $F_{(1.798,16.19)} = 2.75$;<br>p>0.05 | $F_{(1.726,17.26)} = 58.41$;<br>p<0.05 |
| *pre-anesthetic* | 89.1 ± 6.21 | 77.9 ± 12.39 | 74.9 ± 6.96 | 85.0 ± 8.64 | 94.0 ± 8.20 |
| *Induction* | 64.3 ± 11.22* | 107.7 ± 14.79* | 74.1 ± 12.93 | 89.9 ± 9.67 | 60.9 ± 6.36* |
| *Recovery* | 62.4 ± 9.61* | 81.9 ± 18.28 | 81.2 ± 11.90 | 90.3 ± 9.63 | 90.5 ± 8.27 |
| MAP (mm Hg) |  |  |  |  |  |
| F; P Values | $F_{(1.940,21.34)} = 66.89$;<br>p<0.05 | $F_{(1.700,17.00)} = 19.55$;<br>p<0.05 | $F_{(1.409,15.50)} = 2.76$;<br>p>0.05 | $F_{(1.641,14.77)} = 1.82$;<br>p>0.05 | $F_{(1.722,17.22)} = 58.35$;<br>p<0.05 |
| *pre-anesthetic* | 104.4 ± 5.95 | 92.4 ± 11.19 | 93.6 ± 7.33 | 98.6 ± 9.21 | 110.1 ± 9.58 |
| *Induction* | 78.6 ± 11.16* | 127.8 ± 19.71* | 92.3 ± 12.82 | 102.9 ± 9.96 | 74.9 ± 6.80* |
| *Recovery* | 74.6 ± 9.71* | 97.3 ± 20.26 | 98.9 ± 11.67 | 102.9 ± 10.09 | 103.0 ± 8.46 |
| HR (bpm) |  |  |  |  |  |
| F; P Values | $F_{(1.736,19.10)} = 8.22$;<br>p<0.05 | $F_{(1.785,17.85)} = 5.84$;<br>p<0.05 | $F_{(1.771,19.48)} = 72.63$;<br>p<0.05; | $F_{(1.261,11.35)} = 5.31$;<br>p<0.05 | $F_{(1.764,17.64)} = 9.88$;<br>p<0.05 |
| *pre-anesthetic* | 347.4 ± 39.44 | 315.2 ± 42.85 | 372.7 ± 45.35 | 298.1 ± 34.13 | 330.5 ± 40.93 |
| *Induction* | 417.6 ± 42.87* | 272.7 ± 23.37* | 455.6 ± 42.93* | 345.8 ± 26.45* | 355.6 ± 44.40 |
| *Recovery* | 368.7 ± 65.93 | 279.1 ± 60.70 | 468.6 ± 36.36* | 350.8 ± 42.14 | 387.2 ± 48.95* |

Control: Data from pre-anesthesia state in conscious animals, diastolic blood pressure (DBP), mean arterial pressure (MAP), heart rate (HR), systolic blood pressure (SBP), bpm (beats per minute). Data are shown as mean ± SD. One-way ANOVA test for repeated measures, followed by Dunnett's post hoc test. \*p < 0.05 indicates difference compared to the Control value of each group.

Likewise, both SBP, DBP, and MAP were lower ten minutes after anesthesia with ISO (p<0.05), with complete returned to baseline levels during the *recovery* phase (p>0.05). HR was not changed by ISO during *induction* (p>0.05); however, it was elevated in the recovery phase (p<0.05).

For KX we observed a significant increase in baseline SBP, DBP, and MAP (p<0.05) ten minutes after induction, whereas HR was reduced (p<0.05). All hemodynamic parameters returned to baseline levels during the *recovery* phase (p>0.05).

Interestingly, we did not observe any changes in either SBP, DBP, or MAP during *induction* with TBE and TP; however, HR increased (p<0.05) ten minutes thereafter, remaining elevated for TBE (p<0.05) even after recovery of the righting reflex.

Delta (Δ) changes in hemodynamic parameters during the *induction* (SBP: F_(4,51)_= 42.00; p<0.05; DBP: F_(4,51)_= 43.41; p<0.05; MAP: F_(4,51)_= 45.04; p<0.05; HR: F_(4,51)_= 21.78; p<0.05) and *recovery* phases (SBP: F_(4,51)_= 16.49; p<0.05; DBP: F_(4,51)_= 13.85; p<0.05; MAP: F_(4,51)_= 45.04; p<0.05; HR: F_(4,51)_= 9.29; p<0.05) varied significantly among the different anesthetic agents tested (Figure 2).

**Figure 2.**
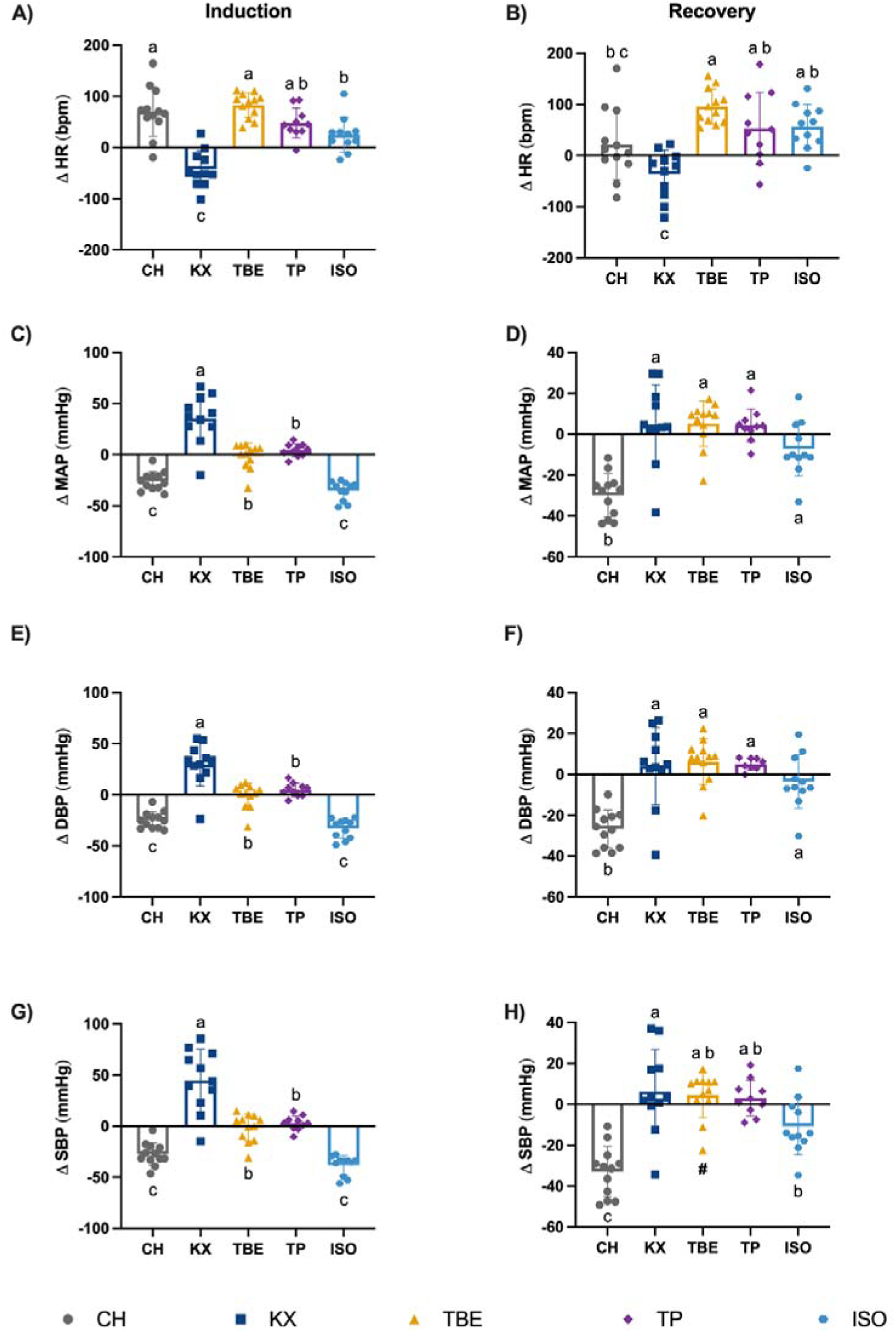
Changes (delta) of the hemodynamic parameters of heart rate (HR, A and B), mean arterial pressure (MAP, C and D), diastolic blood pressure (DBP, E and F) and systolic blood pressure (SBP, G and H) induced by the different anesthetics 10 minutes after induction and in the recovery phase, compared to the pre-anesthetic level. CH: Chloral Hydrate, n = 12, gray; KX: Ketamine and Xylasine, n = 11, dark blue; TBE: Tribromoethanol, n = 12, yellow; TP: Thiopental, n = 10, purple; ISO: Isoflurane, n = 11, light blue. Data are shown as mean ± SD. One-way ANOVA followed by Tukey’s post hoc test. Compact letter display (CLD) indicates significant differences in pairwise comparisons, groups sharing at least one letter do not differ p>0.05. All exact p-values are provided in Table A1 and A2 (Appendix 1 after the References section).

During the *induction* phase, ISO and CH led to reductions in SBP, DBP, and MAP compared to TP, TBE, and the KX (p<0.05; Figure 2 C, E, G; Table A1). In contrast, the KX mixture increased the BP compared to TBE, TP, ISO, and HC (p<0.05). In the recovery phase, SBP, DBP, and MAP remained reduced in the CH group compared to KX, TBE, TP, and ISO (p<0.05) (Figure 2 D, F, H; Table A2). For the ISO group, SBP remained reduced only in comparison to the KX group (p<0.05).

During anesthesia, the KX group significantly reduced HR compared to all other anesthetic agents tested (p<0.05) (Figure 2 A; Table A1). In contrast, CH and TBE significantly increased HR compared to ISO (p<0.05), with no significant differences observed among the remaining anesthetics.

Interestingly, during the *recovery* phase, HR remained significantly reduced in the KX group compared to TBE, TP, and ISO (p<0.05; Figure 2 B; Table A2). In contrast, HR in the TBE group remained elevated relative to CH (p<0.05). No significant differences in HR were observed among other anesthetic agents.

### 3.3. Respiratory Parameters

The effects of anesthesia on respiratory parameters are presented in Table 3. CH and TP reduced V_E_ through reductions in *f_R_* (p<0.05), with no significant effect on V_T_ during the *induction* phase. During *recovery*, both V_E_ and *f_R_*returned to values *pre-anesthesia* for CH. In contrast, *f_R_* remained reduced after recovery in the TP group (p<0.05), though no significant differences were observed for V_E_ (p>0.05).

**Table 3.**
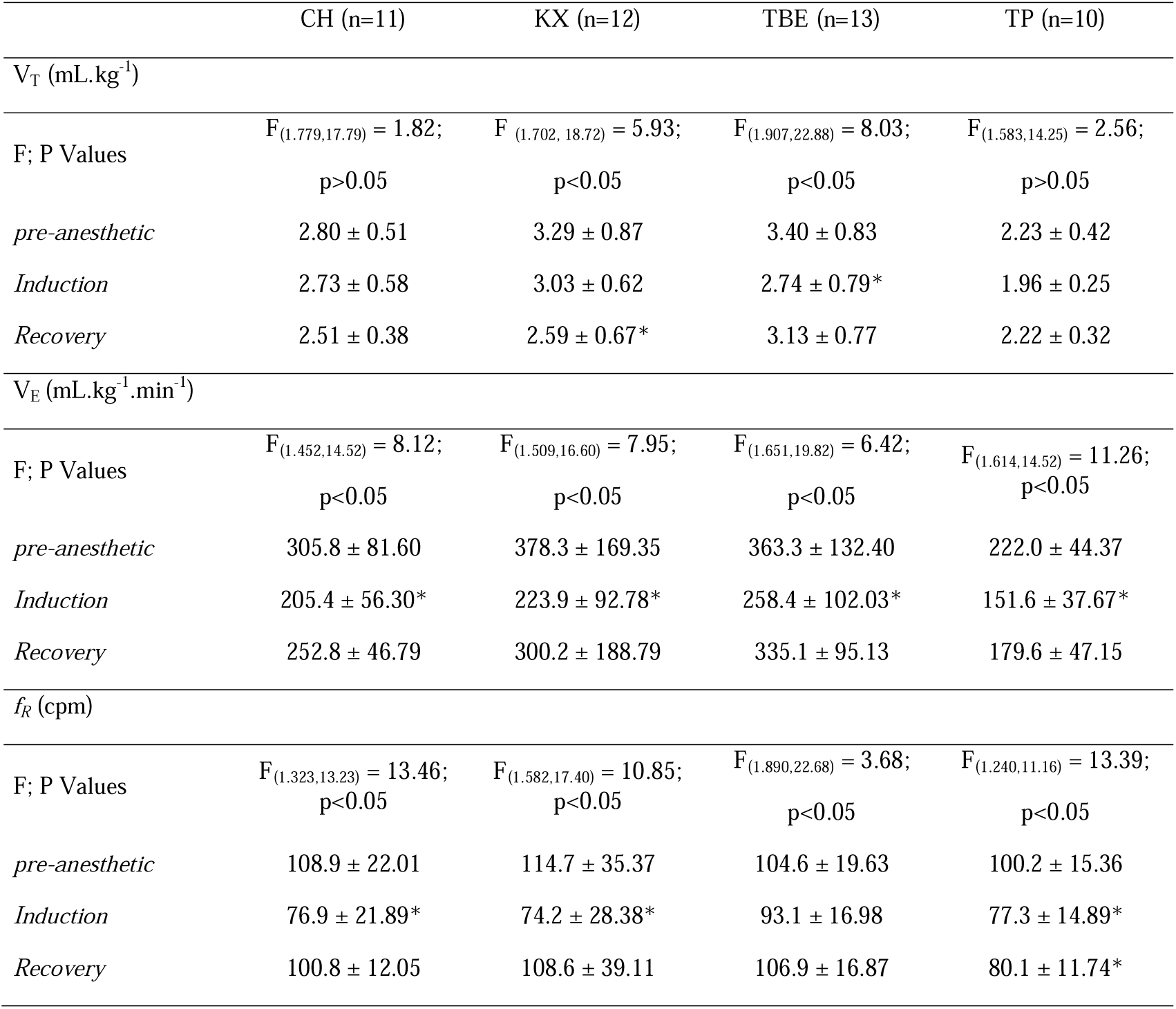
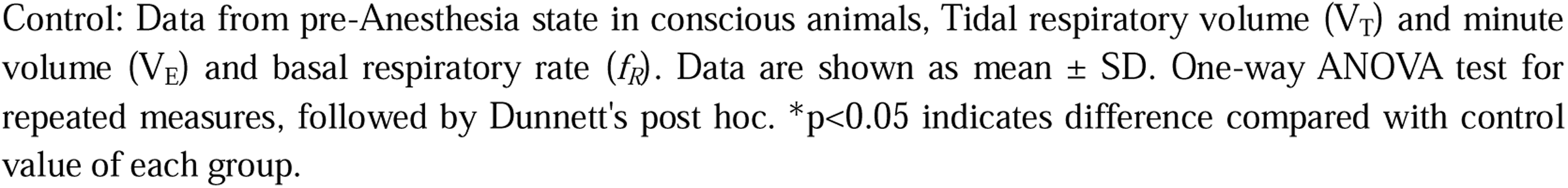
Respiratory data during *pre-anesthetic*, *induction*, and *recovery* phases of anesthesia with Chloral Hydrate (CH), Ketamine/Xylazine (KX), Tribromoethanol (TBE), and Thiopental (TP) in rats.

|  | CH (n=11) | KX (n=12) | TBE (n=13) | TP (n=10) |
| --- | --- | --- | --- | --- |
| $V_T$ (mL.kg<sup>-1</sup>) | | | | |
| F; P Values | $F_{(1.779,17.79)} = 1.82$;<br>$p > 0.05$ | $F_{(1.702, 18.72)} = 5.93$;<br>$p < 0.05$ | $F_{(1.907,22.88)} = 8.03$;<br>$p < 0.05$ | $F_{(1.583,14.25)} = 2.56$;<br>$p > 0.05$ |
| *pre-anesthetic* | $2.80 \pm 0.51$ | $3.29 \pm 0.87$ | $3.40 \pm 0.83$ | $2.23 \pm 0.42$ |
| *Induction* | $2.73 \pm 0.58$ | $3.03 \pm 0.62$ | $2.74 \pm 0.79^*$ | $1.96 \pm 0.25$ |
| *Recovery* | $2.51 \pm 0.38$ | $2.59 \pm 0.67^*$ | $3.13 \pm 0.77$ | $2.22 \pm 0.32$ |
| $V_E$ (mL.kg<sup>-1</sup>.min<sup>-1</sup>) | | | | |
| F; P Values | $F_{(1.452,14.52)} = 8.12$;<br>$p < 0.05$ | $F_{(1.509,16.60)} = 7.95$;<br>$p < 0.05$ | $F_{(1.651,19.82)} = 6.42$;<br>$p < 0.05$ | $F_{(1.614,14.52)} = 11.26$;<br>$p < 0.05$ |
| *pre-anesthetic* | $305.8 \pm 81.60$ | $378.3 \pm 169.35$ | $363.3 \pm 132.40$ | $222.0 \pm 44.37$ |
| *Induction* | $205.4 \pm 56.30^*$ | $223.9 \pm 92.78^*$ | $258.4 \pm 102.03^*$ | $151.6 \pm 37.67^*$ |
| *Recovery* | $252.8 \pm 46.79$ | $300.2 \pm 188.79$ | $335.1 \pm 95.13$ | $179.6 \pm 47.15$ |
| $f_R$ (cpm) | | | | |
| F; P Values | $F_{(1.323,13.23)} = 13.46$;<br>$p < 0.05$ | $F_{(1.582,17.40)} = 10.85$;<br>$p < 0.05$ | $F_{(1.890,22.68)} = 3.68$;<br>$p < 0.05$ | $F_{(1.240,11.16)} = 13.39$;<br>$p < 0.05$ |
| *pre-anesthetic* | $108.9 \pm 22.01$ | $114.7 \pm 35.37$ | $104.6 \pm 19.63$ | $100.2 \pm 15.36$ |
| *Induction* | $76.9 \pm 21.89^*$ | $74.2 \pm 28.38^*$ | $93.1 \pm 16.98$ | $77.3 \pm 14.89^*$ |
| *Recovery* | $100.8 \pm 12.05$ | $108.6 \pm 39.11$ | $106.9 \pm 16.87$ | $80.1 \pm 11.74^*$ |

Control: Data from pre-Anesthesia state in conscious animals, Tidal respiratory volume ( $V_T$ ) and minute volume ( $V_E$ ) and basal respiratory rate ( $f_R$ ). Data are shown as mean ± SD. One-way ANOVA test for repeated measures, followed by Dunnett's post hoc. \* $p < 0.05$ indicates difference compared with control value of each group.

Ten minutes after *induction* with KX, baseline V_E_ and *f_R_*were decreased (p<0.05). Surprisingly, during the *recovery* phase, V_T_ was significantly reduced (p<0.05), whilst both V_E_ and *f_R_* values (p>0.05) were back to *pre-anesthetic* levels.

In contrast to previous anesthetic agents, TBE reduced baseline V_E_ through reductions in V_T_ during the *induction* phase (p<0.05). After recovery of the righting reflex, respiratory parameters returned to values close to *pre-anesthetic* levels (p>0.05).

Among the injectable anesthetics agents, Δ changes during the *induction* phase differed significantly for *f_R_* (F_(3,42)_= 3.39; p<0.05; Figure 3 E; Table A3) but not for V_T_ (F_(3,42)_= 2.03; p>0.05) or V_E_ (F_(3,42)_= 1.33; p>0.05; Figure 3 A and C; Table A3). Although all anesthetics tested numerically reduced V_T_ and V_E_ during *induction*, these reductions were not statistically different between agents (p>0.05). In contrast, KX induced a significantly greater reduction in *f_R_* compared to TBE (p<0.05) (Figure 3 E; Table A3).

**Figure 3.**
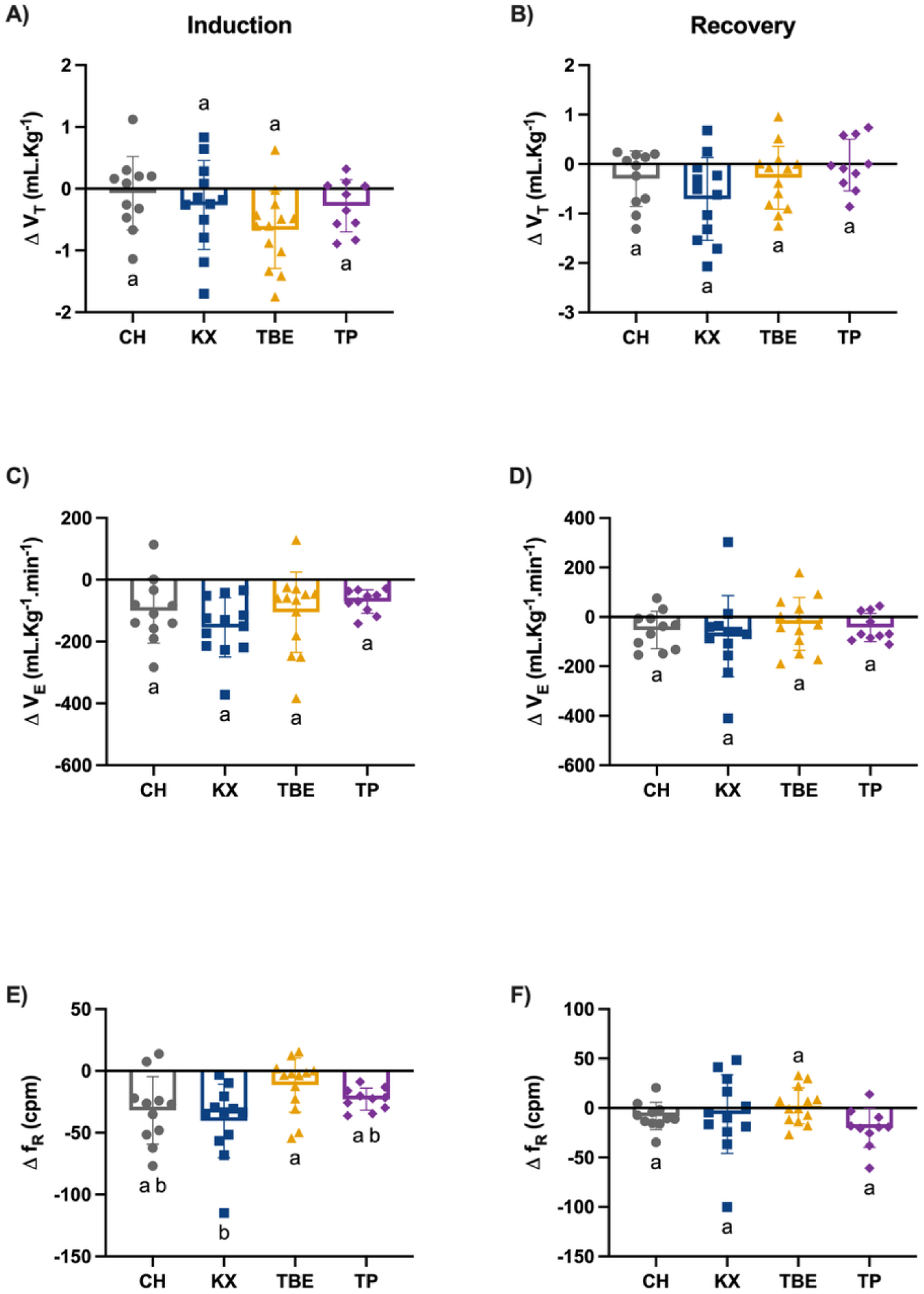
Changes (delta) in tidal respiratory volume (V_T_, A and B), minute volume (V_E_, C and D) and basal respiratory rate (*f*_R_, E and F) induced by the injectable anesthetics 10 minutes after induction and during recovery, compared to the pre-anesthetic level. CH: Chloral Hydrate, n = 11, gray; KX: Ketamine and Xylasine, n = 12, dark blue; TBE: Tribromoethanol, n = 13, yellow; TP: Thiopental, n = 10, purple. Data are shown as mean ± SD. One-way ANOVA followed by Tukey’s post hoc test. Compact letter display (CLD) indicates significant differences in pairwise comparisons, groups sharing at least one letter do not differ p>0.05. All exact p-values are provided in Table A3 and A4 (Appendix 1 after the References section).

During the *recovery* phase, no significant differences were observed in Δ changes among the tested anesthetic agents (V_T_: F_(3,42)_= 2.11; p>0.05; V_E_: F_(3,42)_= 0.44; p>0.05; *f_R_*: F_(3,42)_= 1.48; p>0.05; Figure 3 B, D and F; Table A4).

### 3.4. Time- and frequency-domains cardiovascular variability

The effects of anesthesia on time-domain variability are presented in Table 4. Our data demonstrated anesthetic-specific effects on PI variance, RMSSD and SBP variance. During the *induction* phase, all anesthetic agents reduced PI variance, consistent with our hypothesis. Likewise, SBP variance was significantly reduced for the majority of agents, with the exception of TBE (P>0.05). Additionally, RMSSD was reduced by KX, TBE, and TP.

**Table 4.** Time domain systolic blood pressure (SBP) and pulse interval (PI) variability during *pre-anesthetic*, *induction*, and *recovery* phases of anesthesia with Chloral Hydrate (CH), Ketamine/Xylazine (KX), Tribromoethanol (TBE) and Thiopental (TP) in rats.

|  | CH (n = 11) | KX (n = 9) | TBE (n = 12) | TP (n = 10) | ISO (n = 11) |
| --- | --- | --- | --- | --- | --- |
| **PI Variability** |  |  |  |  |  |
| PI (ms) |  |  |  |  |  |
| F; P Values | F<sub>(1.646, 16.46)</sub> =<br>27.13<br>p<0.05 | F<sub>(1.304, 10.43)</sub> =<br>2.297<br>p>0.05 | F<sub>(1.615, 17.77)</sub> =<br>54.42<br>p<0.05 | F<sub>(1.495, 13.45)</sub> =<br>15.90<br>p<0.05 | F<sub>(1.741, 17.41)</sub> =<br>8.804<br>p<0.05 |
| *pre-anesthetic* | 177.7 ± 20.27 | 196.9 ± 21.32 | 171.5 ± 17.79 | 215.0 ± 18.82 | 188.1 ± 21.85 |
| *Induction* | 148.0 ± 14.73** | 214.6 ± 14.30** | 139.6 ± 9.26** | 173.6 ± 14.43** | 170.8 ± 21.70** |
| *Recovery* | 181.9 ± 17.50 | 212.0 ± 37.77 | 135.2 ± 8.15** | 176.5 ± 16.71** | 166.8 ± 19.01** |
| PI variance (ms<sup>2</sup>) |  |  |  |  |  |
| F; P Values | F<sub>(1.883, 18.83)</sub> =<br>40.32<br>p<0.05 | F<sub>(1.617, 11.32)</sub> =<br>14.92<br>p<0.05 | F<sub>(1.268, 13.95)</sub> =<br>100.6<br>p<0.05 | F<sub>(1.330, 11.97)</sub> =<br>7.563<br>p<0.05 | F<sub>(1.423, 14.23)</sub> =<br>13.44<br>p<0.05 |
| *pre-anesthetic* | 12.0 ± 2.59 | 13.4 ± 3.41 | 13.0 ± 3.37 | 15.6 ± 4.74 | 13.2 ± 3.15 |
| *Induction* | 4.2 ± 1.84** | 6.5 ± 1.77** | 2.6 ± 1.26** | 7.0 ± 3.08** | 5.8 ± 1.33** |
| *Recovery* | 8.8 ± 2.82** | 15.6 ± 5.01 | 3.0 ± 1.02** | 13.2 ± 5.92 | 11.0 ± 5.03 |
| RMSSD (ms) |  |  |  |  |  |
| F; P Values | F<sub>(1.159, 11.59)</sub> =<br>10.76<br>p<0.05 | F<sub>(1.465, 11.72)</sub> =<br>6.439<br>p<0.05 | F<sub>(1.191, 13.10)</sub> =<br>33.92<br>p<0.05 | F<sub>(1.444, 13.00)</sub> =<br>2.290<br>p>0.05 | F<sub>(1.316, 13.16)</sub> =<br>1.370<br>p>0.05 |
| *pre-anesthetic* | 3.1 ± 0.65 | 4.7 ± 1.04 | 3.5 ± 1.13 | 5.1 ± 1.14 | 4.1 ± 1.17 |
| *Induction* | 2.9 ± 0.72 | 3.5 ± 0.47** | 1.6 ± 0.41** | 3.9 ± 1.09* | 3.5 ± 0.27 |
| *Recovery* | 3.9 ± 0.86** | 5.2 ± 1.52 | 1.9 ± 0.56** | 4.5 ± 1.22 | 3.9 ± 1.06 |
| **SBP variability** |  |  |  |  |  |
| SBP variance (mmHg<sup>2</sup>) |  |  |  |  |  |
| F; P Values | F<sub>(1.472, 14.72)</sub> =<br>4.691<br>p<0.05 | F<sub>(1.945, 15.56)</sub> =<br>3.806<br>p<0.05 | F<sub>(1.507, 16.57)</sub> =<br>11.10<br>p<0.05 | F<sub>(1.197, 10.78)</sub> =<br>4.340<br>p>0.05 | F<sub>(1.390, 13.90)</sub> =<br>8.568<br>p<0.05 |
| *pre-anesthetic* | 9.1 ± 2.30 | 8.5 ± 3.00 | 8.1 ± 2.64 | 6.7 ± 1.97 | 7.7 ± 3.10 |
| *Induction* | 5.8 ± 1.98* | 5.2 ± 2.74* | 6.9 ± 2.95 | 4.9 ± 1.71* | 6.2 ± 4.76** |
| *Recovery* | 9.9 ± 5.31 | 8.8 ± 2.60 | 13.9 ± 6.32* | 9.4 ± 5.44 | 10.6 ± 3.27 |

Parameters: Pulse interval (PI) and systolic blood pressure (SBP), Root mean square of successive differences of inter-beat intervals (RMSSD). Data are shown as mean ± SD. One-way ANOVA test for repeated measures, followed by Dunnet's post hoc. \* p<0.05 and \*\*p<0.01 vs pre-anesthetic; \*marginal p-value (0.05 < P < 0.08) vs pre-anesthetic.

During *recovery*, PI variance returned to *pre-anesthetic* levels for most agents, except for CH and TBE. Although SBP variance was not significantly reduced by TBE during *induction*, it was elevated during *recovery*. RMSSD responses varied, with CH inducing a significant increase relative to baseline, TBE maintaining reduced levels, and both KX, TP and ISO returning to *pre-anesthetic* values.

Spectral analysis of HRV in the frequency domain showed that, during the *induction* phase, all anesthetic agents reduced both the VLF and LF bands, including LF/HF ratio (p<0,05), except for ISO, which reached only marginal significance in the LF/HF ratio during *induction* (Table 5). Conversely, the HF bands were increased by all agents during the *induction* phase (p<0.05, Table 5).

**Table 5:**
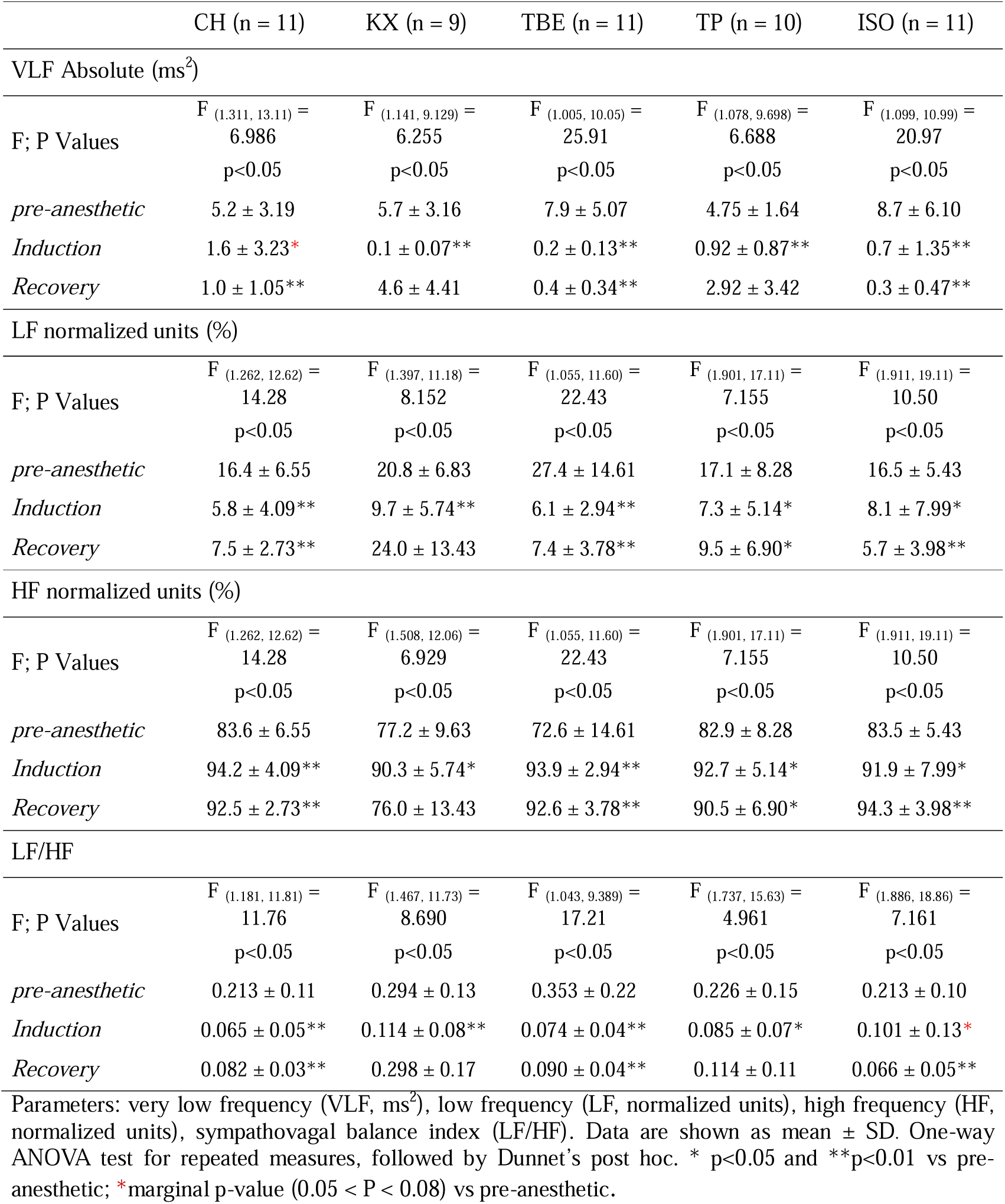
Spectral analysis of pulse intervals (i.e., heart rate) during *pre-anesthetic*, *induction*, and *recovery* phases of anesthesia with Chloral Hydrate (CH), Ketamine/Xylazine (KX), Tribromoethanol (TBE) and Thiopental (TP) in rats.

|  | CH (n = 11) | KX (n = 9) | TBE (n = 11) | TP (n = 10) | ISO (n = 11) |
| --- | --- | --- | --- | --- | --- |
| VLF Absolute (ms<sup>2</sup>) |  |  |  |  |  |
| F; P Values | F<sub>(1.311, 13.11)</sub> =<br>6.986<br>p<0.05 | F<sub>(1.141, 9.129)</sub> =<br>6.255<br>p<0.05 | F<sub>(1.005, 10.05)</sub> =<br>25.91<br>p<0.05 | F<sub>(1.078, 9.698)</sub> =<br>6.688<br>p<0.05 | F<sub>(1.099, 10.99)</sub> =<br>20.97<br>p<0.05 |
| *pre-anesthetic* | 5.2 ± 3.19 | 5.7 ± 3.16 | 7.9 ± 5.07 | 4.75 ± 1.64 | 8.7 ± 6.10 |
| *Induction* | 1.6 ± 3.23* | 0.1 ± 0.07** | 0.2 ± 0.13** | 0.92 ± 0.87** | 0.7 ± 1.35** |
| *Recovery* | 1.0 ± 1.05** | 4.6 ± 4.41 | 0.4 ± 0.34** | 2.92 ± 3.42 | 0.3 ± 0.47** |
| LF normalized units (%) |  |  |  |  |  |
| F; P Values | F<sub>(1.262, 12.62)</sub> =<br>14.28<br>p<0.05 | F<sub>(1.397, 11.18)</sub> =<br>8.152<br>p<0.05 | F<sub>(1.055, 11.60)</sub> =<br>22.43<br>p<0.05 | F<sub>(1.901, 17.11)</sub> =<br>7.155<br>p<0.05 | F<sub>(1.911, 19.11)</sub> =<br>10.50<br>p<0.05 |
| *pre-anesthetic* | 16.4 ± 6.55 | 20.8 ± 6.83 | 27.4 ± 14.61 | 17.1 ± 8.28 | 16.5 ± 5.43 |
| *Induction* | 5.8 ± 4.09** | 9.7 ± 5.74** | 6.1 ± 2.94** | 7.3 ± 5.14* | 8.1 ± 7.99* |
| *Recovery* | 7.5 ± 2.73** | 24.0 ± 13.43 | 7.4 ± 3.78** | 9.5 ± 6.90* | 5.7 ± 3.98** |
| HF normalized units (%) |  |  |  |  |  |
| F; P Values | F<sub>(1.262, 12.62)</sub> =<br>14.28<br>p<0.05 | F<sub>(1.508, 12.06)</sub> =<br>6.929<br>p<0.05 | F<sub>(1.055, 11.60)</sub> =<br>22.43<br>p<0.05 | F<sub>(1.901, 17.11)</sub> =<br>7.155<br>p<0.05 | F<sub>(1.911, 19.11)</sub> =<br>10.50<br>p<0.05 |
| *pre-anesthetic* | 83.6 ± 6.55 | 77.2 ± 9.63 | 72.6 ± 14.61 | 82.9 ± 8.28 | 83.5 ± 5.43 |
| *Induction* | 94.2 ± 4.09** | 90.3 ± 5.74* | 93.9 ± 2.94** | 92.7 ± 5.14* | 91.9 ± 7.99* |
| *Recovery* | 92.5 ± 2.73** | 76.0 ± 13.43 | 92.6 ± 3.78** | 90.5 ± 6.90* | 94.3 ± 3.98** |
| LF/HF |  |  |  |  |  |
| F; P Values | F<sub>(1.181, 11.81)</sub> =<br>11.76<br>p<0.05 | F<sub>(1.467, 11.73)</sub> =<br>8.690<br>p<0.05 | F<sub>(1.043, 9.389)</sub> =<br>17.21<br>p<0.05 | F<sub>(1.737, 15.63)</sub> =<br>4.961<br>p<0.05 | F<sub>(1.886, 18.86)</sub> =<br>7.161<br>p<0.05 |
| *pre-anesthetic* | 0.213 ± 0.11 | 0.294 ± 0.13 | 0.353 ± 0.22 | 0.226 ± 0.15 | 0.213 ± 0.10 |
| *Induction* | 0.065 ± 0.05** | 0.114 ± 0.08** | 0.074 ± 0.04** | 0.085 ± 0.07* | 0.101 ± 0.13* |
| *Recovery* | 0.082 ± 0.03** | 0.298 ± 0.17 | 0.090 ± 0.04** | 0.114 ± 0.11 | 0.066 ± 0.05** |

Parameters: very low frequency (VLF, ms<sup>2</sup>), low frequency (LF, normalized units), high frequency (HF, normalized units), sympathovagal balance index (LF/HF). Data are shown as mean ± SD. One-way ANOVA test for repeated measures, followed by Dunnet's post hoc. \* p<0.05 and \*\*p<0.01 vs pre-anesthetic; \*marginal p-value (0.05 < P < 0.08) vs pre-anesthetic.

During *recovery*, VLF returned to *pre-anesthetic* levels for KX and TP but remained reduced for CH, TBE and ISO. LF bands remained reduced, and HF increased during *recovery* for all anesthetic agents, except for KX (Table 5). LF/HF ratio was not statistically different for KX and TP during *recovery* phase (p>0.05).

The effects of anesthesia on baroreceptors were assessed via analysis of sBRS. Data representing the gain for up-sequences, down-sequences, all-sequences and BEI are depicted in Table 6.

**Table 6.**
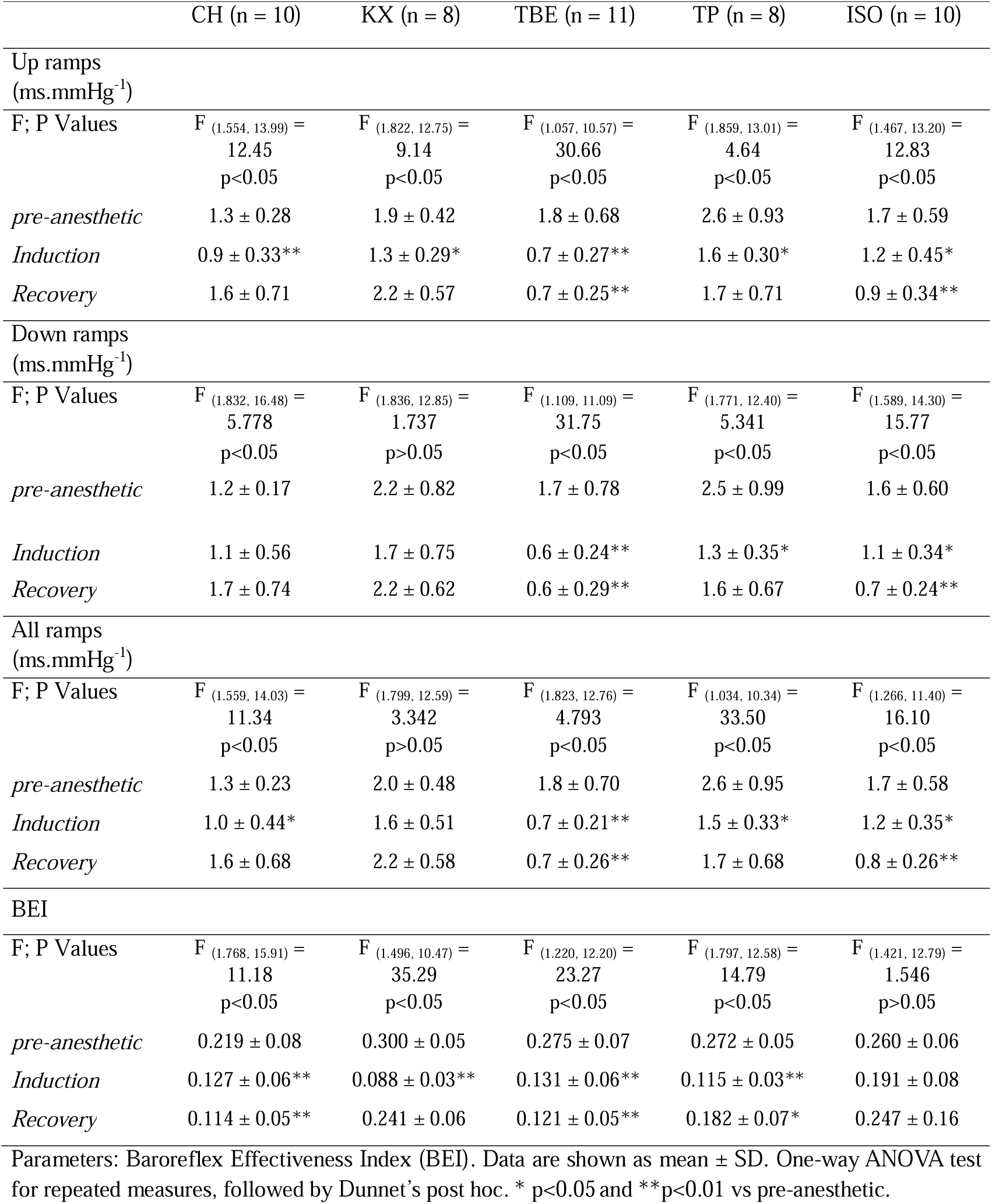
Spontaneous baroreflex data of during *pre-anesthetic, induction, and recovery phases* of anesthesia with Chloral Hydrate (CH), Ketamine/Xylazine (KX), Tribromoethanol (TBE) and Thiopental (TP) in rats.

During induction, all anesthetic agents significantly reduced the gain for up-ramp sequences, as well as the BEI (p<0.05; Table 6), except for ISO. The gains for down-ramps were significantly reduced during the *induction* phase only with TBE, TP and ISO and all-ramps with CH, TBE, TP and ISO (p<0.05).

During *recovery*, the gains for up-ramps and all-ramps returned to *pre-anesthetic* levels for CH, KX and TP, while down-ramps remained reduced for TBE and ISO. The BEI stayed reduced during the *recovery* phase with CH, TBE and TP (Table 6).

Δ changes in BEI varied significantly among the different anesthetic agents during the *induction* phase (F_(4,42)_= 4.077; p<0.01), but not during the *recovery* (F_(4,42)_= 2.220; p>0.05) (Figure 4; Table A5).

**Figure 4.**
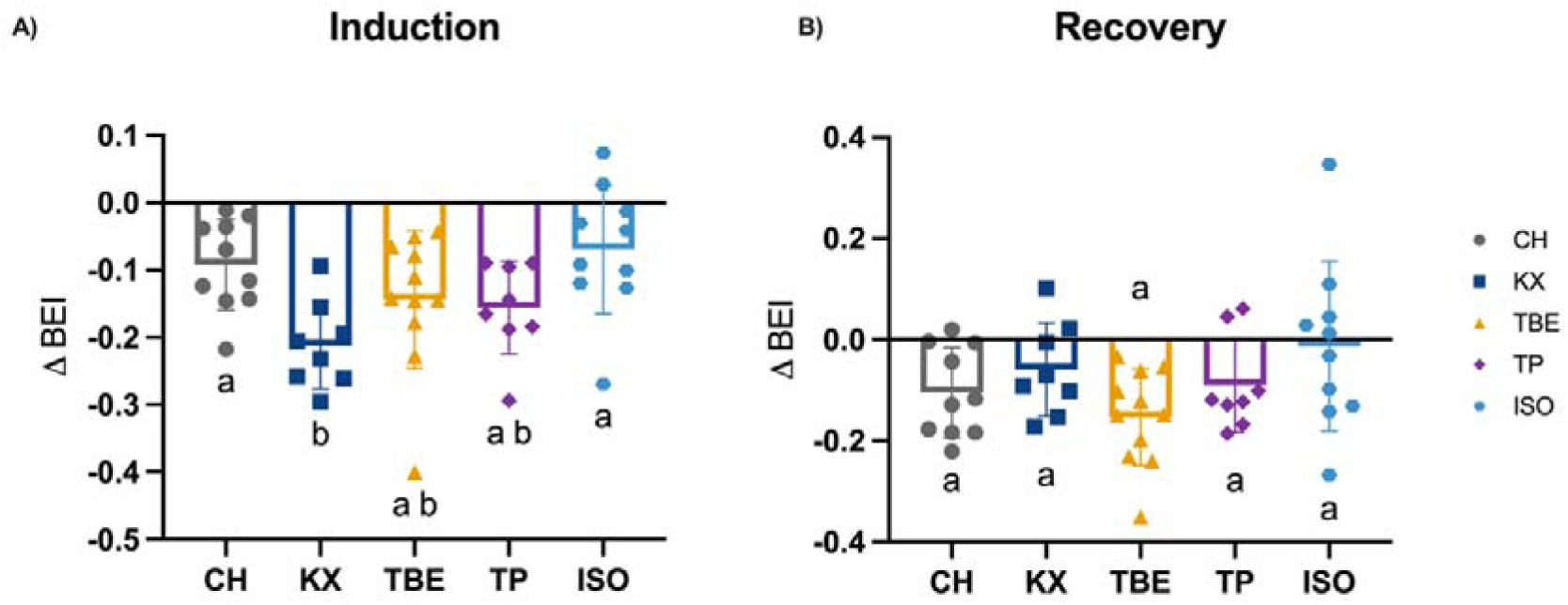
Changes in Baroreflex Effectiveness Index (BEI) during induction (panel A) and recovery (panel B) phases of anesthesia with chloral hydrate (CH, n = 10, gray), ketamine/xylazine (KX, n = 8, dark blue), tribromoethanol (TBE, n = 11, yellow), thiopental (TP, n = 8, purple) and isoflurane (ISO, n = 10, light blue) in rats. Data are shown as mean ± SD. One-way ANOVA followed by Tukey’s post hoc test. Compact letter display (CLD) indicates significant differences in pairwise comparisons, groups sharing at least one letter do not differ p>0.05. All exact p-values are provided in Table A5 (Appendix 1 after the References section).

During *induction*, KX produced the greatest reduction in BEI among all anesthetic agents, with statistically significant differences compared to CH and ISO. However, these differences were no longer evident during the *recovery* phase (Figure 4; Table A5).

### 3.5. Biochemical Parameters

#### 3.5.1. Effect of Anesthesia on Blood Gases

The effect of the different anesthetics on blood gas parameters *pre* and *post-induction* is shown in Table 7. The CH reduced the pH, HCO_3_, and total CO_2_ levels (p<0.05), with no statistically significant change being observed for the other parameters analyzed.

**Table 7.**
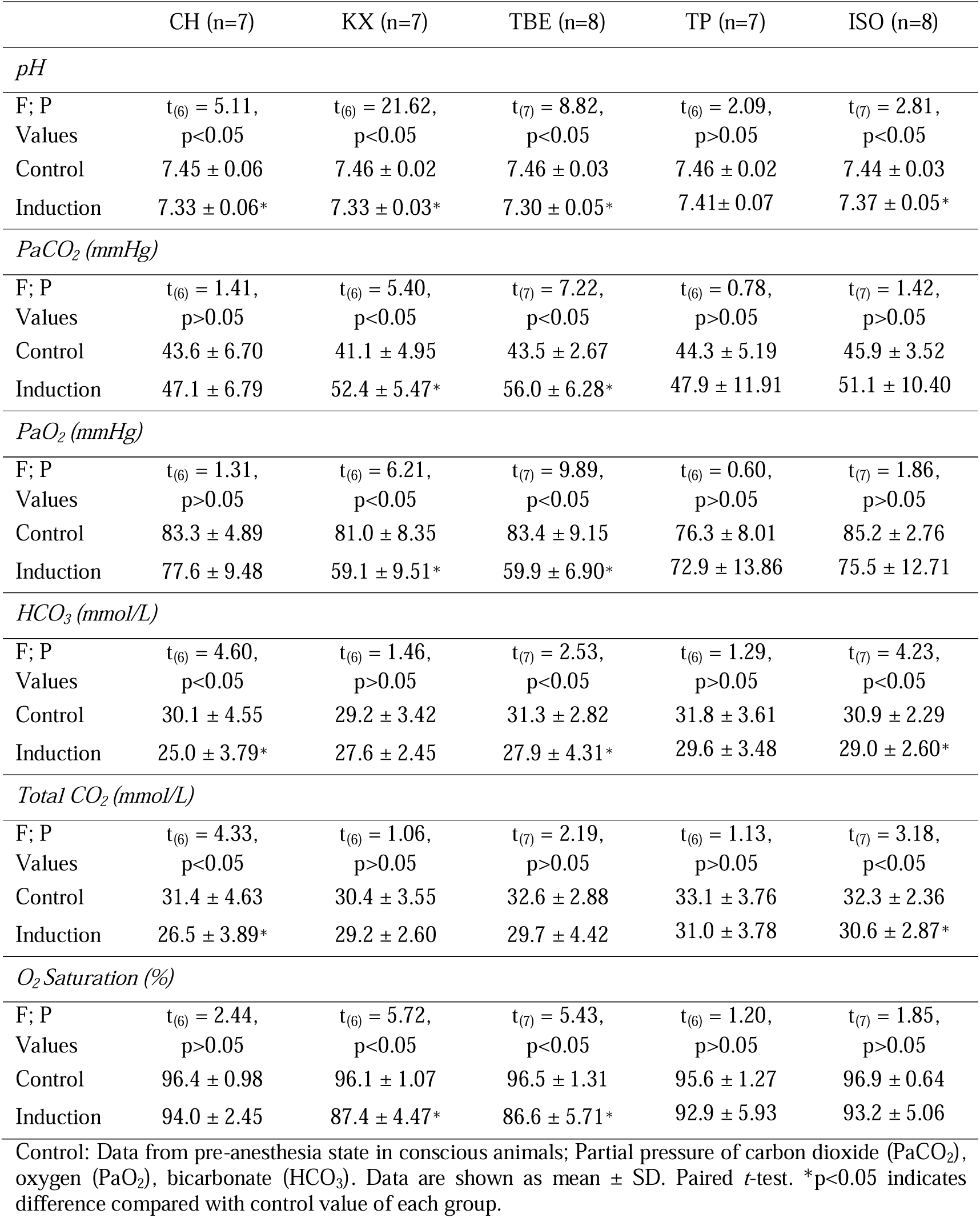
Blood gas data before and after 10 minutes of *induction* of anesthesia with Chloral Hydrate (CH), Ketamine/Xylazine (KX), Tribromoethanol (TBE), Thiopental (TP) and Isoflurane (ISO) in rats.

The KX mixture reduced the pH, PaO_2_, and O_2_ saturation (p<0.05) and, in contrast, increased the PaCO_2_ (p<0.05), with no statistically significant change in the HCO_3_, and total CO_2_ levels compared to baseline.

*Induction* with TBE reduced the pH, PaO_2_, O_2_ saturation, and HCO_3_ levels (p<0.05) and increased the PaCO_2_ (p<0.05), with no change in total CO_2_.

The pH, PaCO_2_, and HCO_3_ (p<0.05) were reduced after induction with ISO, with no change in the other parameters analyzed. Induction with TP did not change all parameters analyzed.

Anesthetics changed the pH, PaO_2_ and O_2_ saturation differently (pH: F_(4, 32)_ = 4.18; p<0.05; PaO_2_: F_(4, 32)_ = 4.45; p<0.05; O_2_ saturation: F_(4, 32)_ = 3.97; p<0.05, Figure 5 A, C, F; Table A6), but not the PaCO_2,_ HCO_3_ and Total CO_2_ (pCO_2_: F_(4, 32)_ = 1.97; p>0.05; HCO_3_: F_(4, 32)_ = 1.39; p>0.05; Total CO_2_: F_(4, 32)_ = 1.34; p>0.05; Figure 5 B, D, E; Table A6). TBE induced a greater reduction in pH compared to TP and ISO (p<0.05), with no difference among the other anesthetics tested. The KX mixture led to the smallest reduction in pO_2_ compared to TBE and TP (p<0.05), with no difference among the other anesthetics tested. CH induced a smaller reduction in O_2_ saturation when compared to TBE (p<0.05). All anesthetics reduced HCO_3_ and total CO_2_ levels and increased PaCO_2_, with no statistically significant differences being observed among the anesthetics (Figure 5 B, D, E; Table A6).

**Figure 5.**
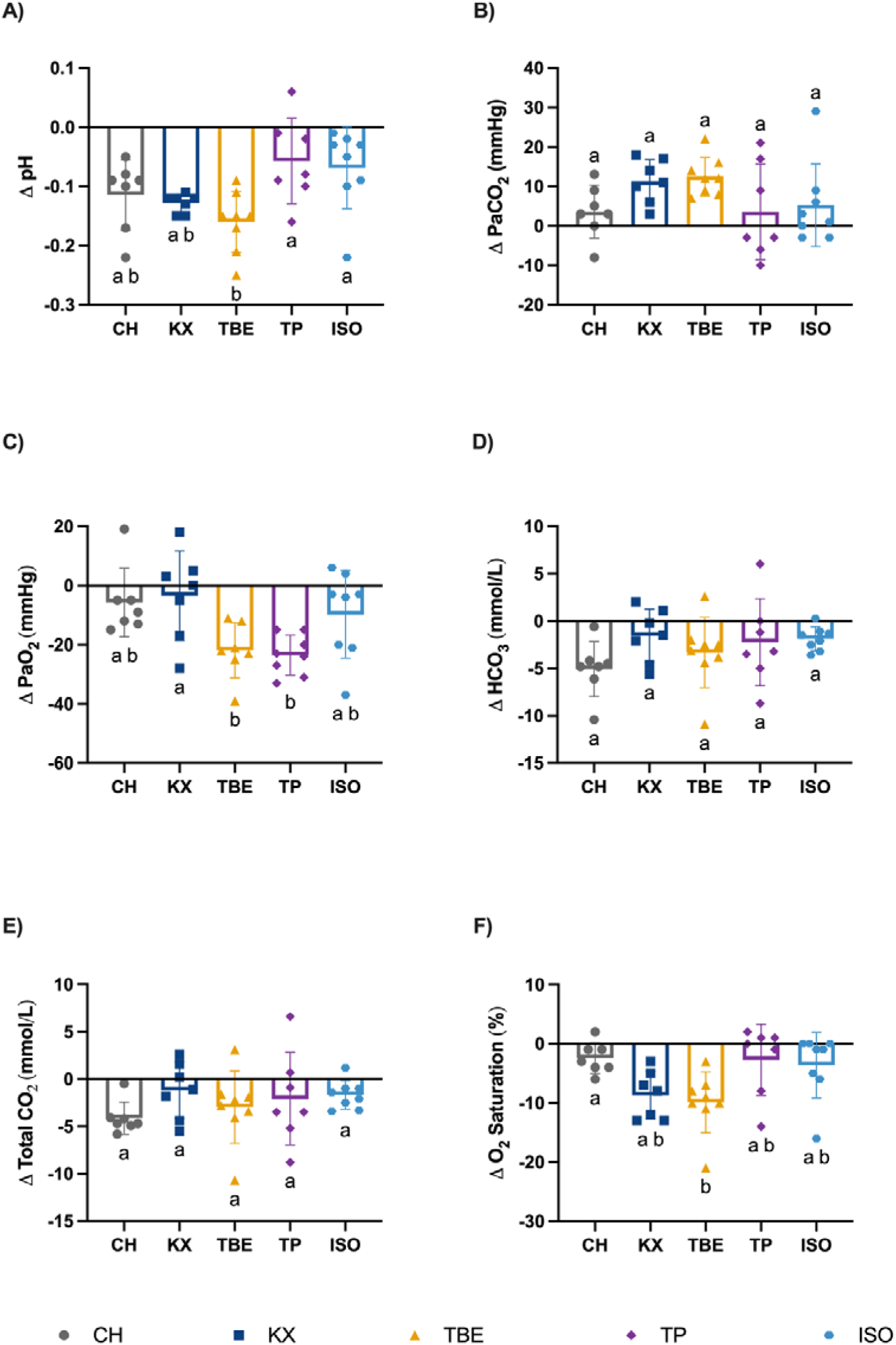
Changes (delta) induced by the different anesthetics in pH (A), PaCO_2_ (B), PaO_2_ (C), HCO_3_ (D), Total CO_2_ (E) and O_2_ saturation (F) across experimental groups: chloral hydrate (CH, n = 7, gray), ketamine/xylazine (KX, n = 7, dark blue), tribromoethanol (TBE, n = 8, yellow), thiopental (TP, n = 7, purple) and isoflurane (ISO, n = 8, light blue) in rats. Data are shown as mean ± SD. One-way ANOVA followed by Tukey’s post hoc test. Compact letter display (CLD) indicates significant differences in pairwise comparisons, groups sharing at least one letter do not differ p>0.05. All exact p-values are provided in Table A6 (Appendix 1 after the References section).

#### 3.5.2. Anesthesia effects on cardiac biomarkers

The effect of different anesthetics on the butyrylcholinesterase (BChE), Troponin I (cTnI), Lactate dehydrogenase (LDH), and creatine kinase MB (CK-MB) during the induction phase is shown in Table 8.

**Table 8.** Cardiac biomarkers measured before and after 10 minutes of *induction* of anesthesia with Chloral Hydrate (CH), Ketamine/Xylazine (KX), Tribromoethanol (TBE), Thiopental (TP) and Isoflurane (ISO) in rats.

|  | CH (n=8) | KX (n=8) | TBE (n=8) | TP (n=8) | ISO (n=8) |
| --- | --- | --- | --- | --- | --- |
| *BChE* (U/L) |  |  |  |  |  |
| F; P Values | $t_{(7)} = 2.00$,<br>$p > 0.05$ | $t_{(7)} = 7.00$,<br>$p < 0.05$ | $t_{(6)} = 3.29$,<br>$p < 0.05$ | $t_{(7)} = 0.00$,<br>$p > 0.05$ | $t_{(7)} = 2.64$,<br>$p < 0.05$ |
| Control | $18.4 \pm 3.62$ | $23.6 \pm 6.18$ | $17.5 \pm 2.86$ | $17.1 \pm 3.47$ | $18.6 \pm 5.85$ |
| Induction | $14.9 \pm 4.08$ | $13.1 \pm 2.48^*$ | $14.5 \pm 1.32^*$ | $17.1 \pm 2.92$ | $15.2 \pm 5.03^*$ |
| *cTnI* (ng/L) |  |  |  |  |  |
| F; P Values | $t_{(7)} = 0.16$,<br>$p > 0.05$ | $t_{(5)} = 2.35$,<br>$p > 0.05$ | $t_{(6)} = 0.77$,<br>$p > 0.05$ | $t_{(7)} = 0.83$,<br>$p > 0.05$ | $t_{(7)} = 0.94$,<br>$p > 0.05$ |
| Control | $1.97 \pm 2.64$ | $1.26 \pm 0.79$ | $1.34 \pm 0.41$ | $2.9 \pm 3.28$ | $1.61 \pm 1.46$ |
| Induction | $2.05 \pm 1.63$ | $0.81 \pm 0.57$ | $1.23 \pm 0.16$ | $3.1 \pm 3.68$ | $1.25 \pm 1.42$ |
| *LDH* (U/L) |  |  |  |  |  |
| F; P Values | $t_{(7)} = 1.28$,<br>$p > 0.05$ | $t_{(7)} = 1.11$,<br>$p > 0.05$ | $t_{(7)} = 1.12$,<br>$p > 0.05$ | $t_{(7)} = 0.69$,<br>$p > 0.05$ | $t_{(7)} = 1.27$,<br>$p > 0.05$ |
| Control | $334.3 \pm 135.0$ | $237.4 \pm 141.9$ | $408.5 \pm 76.89$ | $268.0 \pm 188.8$ | $251.4 \pm 51.20$ |
| Induction | $389.6 \pm 109.6$ | $254.1 \pm 161.9$ | $379.1 \pm 118.4$ | $252.5 \pm 183.7$ | $282.6 \pm 79.99$ |
| *CK-MB* (U/L) |  |  |  |  |  |
| F; P Values | $t_{(4)} = 3.45$,<br>$p < 0.05$ | $t_{(7)} = 3.26$,<br>$p < 0.05$ | $t_{(7)} = 1.28$,<br>$p > 0.05$ | $t_{(7)} = 0.09$,<br>$p > 0.05$ | $t_{(7)} = 1.77$,<br>$p > 0.05$ |
| Control | $306.6 \pm 62.87$ | $263.0 \pm 66.42$ | $334.3 \pm 135.0$ | $270.8 \pm 79.16$ | $300.4 \pm 47.74$ |
| Induction | $421.4 \pm 130.5^*$ | $325.6 \pm 72.51^*$ | $389.6 \pm 109.6$ | $268.4 \pm 61.88$ | $349.8 \pm 88.12$ |

Control: Data from pre-anesthesia state in conscious animals, butyrylcholinesterase enzyme (BChE), Troponin I (cTnI), Lactate dehydrogenase (LDH) and creatine kinase MB fraction (CK-MB). Data are shown as mean ± SD. Paired *t*-test. \* $p < 0.05$ indicates difference compared to control value of each group.

CH increased CK-MB (p<0.05), without changing the other biomarkers tested. The KX mixture reduced the BChE enzyme activity (p<0.05) and increased the CK-MB (p<0.05), with the remaining biomarkers being unchanged. Similarly, TBE and ISO reduced the BChE activity (p<0.05), but without significant change on the other proteins. The four proteins analyzed remained unchanged by TP during the induction phase of anesthesia (p>0.05).

Anesthetics changed differently the CK-MB and BChE activities (CK-MB: F_(4, 34)_ = 3.63; p<0.05; BChE: F_(4, 34)_ = 9.06, p<0.05, Figure 6 A and C; Table A7), but not the LDH and TnI (LDH: F_(4, 35)_ = 2.23; p>0.05; cTnI: F_(4, 31)_ = 0.70, p>0.05, Figure 5 B and C; Table A7). KX mixture induced a greater reduction in BChE activity compared to CH, TBE, TP, and ISO (p<0.05), with no difference observed among the other anesthetics tested. CH increased the CK-MB (p<0.05) compared with TP and TBE, with no difference observed between the other anesthetics tested.

**Figure 6.**
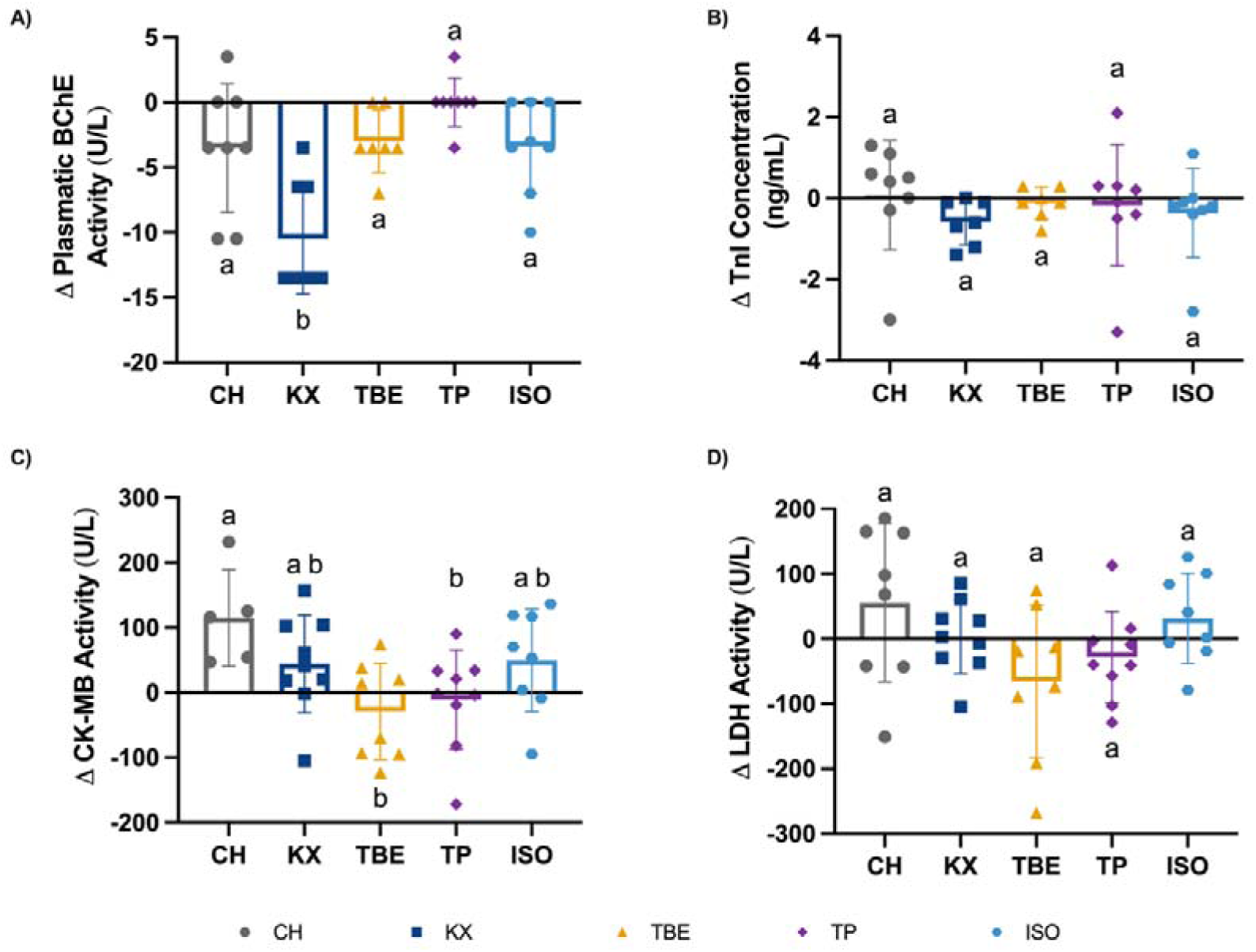
Changes (delta) during the *induction* phase of the enzymes plasmatic butyrylcholinesterase (BChE, A) in (U/L), Troponin I (TnI, B) in (ng/mL), creatine kinase MB fraction (CK-MB, C) (U/L) and Lactate dehydrogenase (LDH, D) in (U/L) induced by the different anesthetics tested: chloral hydrate (CH, gray), ketamine/xylazine (KX, dark blue), tribromoethanol (TBE, yellow), thiopental (TP, purple) and isoflurane (ISO, light blue) in rats. Data are shown as mean ± SD. One-way ANOVA followed by Tukey’s post hoc test. Compact letter display (CLD) indicates significant differences in pairwise comparisons, groups sharing at least one letter do not differ p>0.05. All exact p-values are provided in Table A7 (Appendix 1 after the References section).

## 4. Discussion

Our findings demonstrate significant differences in how commonly used anesthetic agents influence cardiorespiratory parameters and cardiac biomarkers in preclinical models. Among the agents tested, TBE exhibited the fastest onset of hypnosis and the quickest recovery of consciousness, suggesting a rapid pharmacodynamic profile. In contrast, TP required the longest time to induce loss of consciousness, as well as delayed loss of pedal reflex and tail pinch response, followed closely by CH. Hemodynamic data revealed that both CH and ISO caused reductions in blood pressure, with CH showing a prolonged hypotensive effect during recovery. An increase in HR during CH induction suggests preserved baroreceptor function, whereas ISO failed to elicit a compensatory tachycardia, indicating possible baroreflex attenuation. Interestingly, KX induced a rise in blood pressure during anesthesia, followed by a baroreceptor-mediated decrease in HR. TBE and TP, on the other hand, led to transient increases in HR during induction only. Regarding cardiac biomarkers, nearly all anesthetic agents reduced BChE activity, except TP and CH. Although no statistical differences were observed in cTnI and LDH, both CH and KX elicited elevations in CK-MB, suggesting potential myocardial stress or a direct interaction between the anesthetic agents and either the enzyme or reagents used in the assay.

### 4.1. Onset and Depth of Anesthesia

During the *induction* phase, we observed variable onset times of hypnosis among the different anesthetic agents. As previously noted, TBE exhibited the fastest onset of hypnosis and the quickest recovery of consciousness, with loss of consciousness occurring approximately 69 seconds post-administration, followed by loss of pedal reflex and tail pinch at around 102 seconds. The total duration of anesthesia was approximately 15 minutes. These findings are consistent with previous reports in the literature(Meyer & Fish, 2005b).

In contrast, TP and CH required significantly longer times to induce hypnosis and analgesia. Abdi-Azar and Maleki (2014) reported that intraperitoneal injections of TP required nearly five minutes to achieve complete loss of consciousness and analgesia. However, Goldstein and Aronow (1959) demonstrated much faster onset, with loss of the righting reflex occurring within 30 seconds following intravenous administration of TP at 15 mg/kg.

Field et al. (1993) described the sequence of anesthetic effects in rodents administered CH, beginning with loss of muscle tone and palpebral reflex, followed by loss of response to tail pinch, and culminating in the disappearance of the corneal reflex. They reported an induction period of less than three minutes and a mean anesthetic duration of 111 minutes at a dose of 400 mg/kg (i.p.), which aligns with our observations. Similarly, Sisson and Siegel (1989) reported an anesthetic duration of approximately 60 minutes with CH. More recently, Shcherbak et al. (2024) found that a single intraperitoneal dose of CH at 400 mg/kg in aged Wistar rats (24 months) induced anesthesia but was associated with a high mortality rate (37.5%) within 48 hours, highlighting the increased vulnerability of older animals to this agent.

### 4.2. Cardiovascular Effects of Anesthesia

Our data indicate that different anesthetic agents exert distinct effects on hemodynamic parameters. Both CH and ISO significantly reduced BP, specifically MAP, SBP, and DBP, during the *induction* phase. Notably, CH anesthesia elicited a compensatory tachycardia, suggesting preserved baroreflex function. This response was absent with ISO. This interpretation is supported by our analysis of sBRS, which showed that the average gain of down-ramp sequences remained unchanged under CH, whereas it was significantly attenuated under ISO.

It is important to note that the gain of baroreflex down-ramp sequences does not result from baroreceptor activation per se, but rather from baroreceptor unloading. When BP falls, the tonic inhibitory input from baroreceptors to pre-motor sympathetic neurons in the rostral ventrolateral medulla (RVLM) is withdrawn. This disinhibition leads to increased peripheral sympathetic outflow, including activation of cardiac sympathetic efferents (Guyenet et al., 2018), which in turn drives the observed tachycardia.

The reason why CH preserves this unloading-mediated baroreflex response while ISO does not remains unclear. Interestingly, both anesthetic agents significantly reduced the gain of up-ramp sequences and induced hypotension, which may suggest a direct inhibitory effect on RVLM neurons. This hypothesis is supported by our spectral analysis of HRV, which shows reductions in both the LF band and the LF/HF ratio across all anesthetic agents.

However, if RVLM inhibition were the sole mechanism, we would expect suppression of both arms of the baroreflex response. This discrepancy leads us to speculate that the CH-induced fall in BP may involve additional actions on other brainstem nuclei, possibly the nucleus tractus solitarius (NTS). Another potential mechanism underlying CH-induced cardiovascular depression may relate to its sedative properties, which resemble those of ethanol. Like ethanol, CH may increase neuronal membrane permeability, potentially leading to suppression of both vascular and cardiac function (Vardanyan & Hruby, 2006).

TP and TBE increased HR without significantly affecting BP, which is consistent with previous reports in rats (Abdi-Azar & Maleki, 2014; Kushawaha et al., 2011). This effect is speculated to result from baroreflex attenuation, which renders the system less capable of suppressing sympathetic outflow (Aono et al., 2001). Our sBRS analysis supports this interpretation, showing significantly reduced gains in both down-and up-ramp sequences. Although spectral analysis indicates a reduction in sympathetic activity, evidenced by decreases in the LF band and LF/HF ratio, we still observed a reduction in the BEI. This suggests an overall decrease in the number of active baroreflex sequences, pointing to impaired baroreflex function despite reduced sympathetic tone.

This reduction may also help explain why KX produced a rise in blood pressure. Although the gain of baroreflex sequences was not markedly affected, i.e., the reduction in average gain for up-ramp sequences was not numerically large, KX exhibited the lowest proportion of baroreflex sequences during the induction phase (8.8%). This suggests that under KX, baroreflex sensitivity remains relatively intact, but the probability of evoking a baroreflex response is diminished.

Additional studies have emphasized the variability in cardiovascular outcomes during KX anesthesia, with transient tachycardic responses occasionally reported. These are likely attributable to the sympathomimetic effects of ketamine and the specific dose ratio with xylazine (Wellington et al., 2013). Such discrepancies underscore the critical influence of drug proportions: ketamine alone is known to increase both BP and HR (Haskins et al., 1985), whereas xylazine counteracts these effects through α_2_-adrenergic inhibition of catecholamine release (Greene & Thurmon, 1988; Sumitra et al., 2004).

### 4.3. Respiratory Effects of Anesthesia

Our findings demonstrate that, regardless of the anesthetic agent used, anesthesia consistently attenuates respiratory function. This attenuation was primarily due to reductions in *f_R_*, leading to a decrease in V_E_. Consequently, we observed a drop in PaO_2_ and oxygen saturation, alongside a rise in PaCO_2_ and a fall in blood pH, indicative of significant respiratory depression.

For CH, both *f_R_* and V_E_ were reduced in our study. Similar findings were reported by Zausinger et al. (2002), who observed respiratory depression in rats with CH. The authors observed hypercapnia and acidosis confirming its depressant effects. TP, KX, and TBE also produced marked but distinct effects on ventilatory function. In our study, TP significantly reduced *f_R_* and V_E_, while V_T_ remained stable. In rats, Wada et al. (1996) demonstrated that TP produced dose-dependent alterations in blood gases, with higher doses leading to sustained elevations in PaCO□ and lower pH. In another comparative study, Sumitra et al. (2004) reported that TP depressed *f_R_*, although the magnitude was less pronounced than with KX. Similarly, our results revealed that KX evoked more profound respiratory dysfunction, with decreases in *f_R_*, V_T_, and V_E_. This conclusion is supported by many studies in the literature (Ajadi et al., 2013; Massey & Richerson, 2017; Schwarzkopf et al., 2013; Sumitra et al., 2004).

ISO also influenced ventilatory control and blood-gas status. In our study, ISO was associated with reduced PaO□ and pH and increased PaCO□, consistent with respiratory depression. In disagreement with our findings, Schwarzkopf et al. (2013) reported that volatile anesthetic agents, including ISO, did not induce respiratory acidosis to the same level as injectable ones, for instance CH and KX. On the other hand, Zausinger et al. (2002) described that volatile agents depressed respiration in rats, including hypercapnia and acidosis.

To explain these conflicting results, Massey and Richerson (2017) demonstrated that low concentrations of ISO (0.5–1%) markedly attenuate central chemoreceptor activity, as evidenced by a reduced slope between V_E_ and inspired CO□ concentration, i.e., a blunted hypercapnic ventilatory response. In their study, KX also depressed baseline V_E_ and attenuated CO□ responses *in vivo*.

The underlying mechanism of reduced central chemosensitivity remains unknown. However, Massey et al. (2015) showed that ISO suppresses the responsiveness of brainstem serotonergic neurons to pH/CO□. In contrast, the blunted CO□ reflex observed with KX *in vivo* does not result from impaired neuronal chemosensitivity, as the excitability of serotonergic chemosensitive neurons remained intact. Taken together, these findings suggest that both ISO and KX impair ventilatory control and blunt chemosensory responses, albeit through distinct mechanisms.

### 4.4. Cardiac biomarkers

Our study also aimed to evaluate whether commonly used anesthetic agents interfere with blood biomarkers associated with cardiac injury. CK-MB, LDH, cTnI, and BChE were measured before and after *induction* to allow within-subject comparisons. Although BChE is not routinely used as a clinical biomarker of myocardial damage, it has gained relevance in cardiovascular research due to its prognostic value for mortality in patients with acute myocardial infarction (Brzezinski-Sinai et al., 2021; Michels et al., 2021; Sun et al., 2016). Serum BChE activity is associated with cardiovascular prognosis, with reduced levels often reflecting systemic inflammation (Zivkovic et al., 2015), poorer cardiac function (Sun et al., 2016), a higher risk of mortality from coronary heart disease (Goliasch et al., 2012) and myocardial infarction (Sun et al., 2016).

Only a few studies have addressed how anesthesia affects BChE activity. Sinai et al. (2021) demonstrated that both injectable and volatile anesthetics reduced cholinergic status during surgery in patients with and without atherosclerosis, by altering acetylcholinesterase and BChE activity. Similarly, Carrasco et al. (1978) reported that ketamine and two volatile agents (e.g., enflurane and halothane) reduced BChE activity intraoperatively compared to preoperative levels. Consistent with these findings, Nana et al. (1977) showed that general anesthesia with TP and halothane significantly reduced BChE activity. Taken together, these studies support our findings and highlight the potential for broader anesthetic interference with BChE activity.

Beyond the BChE activity, the injectable anesthetics agents tested in our study exhibited distinct effects on classical biomarkers of myocardial injury. CH and KX were associated with elevations in CK-MB, whereas LDH and cTnI remained stable across groups. In contrast, TP demonstrated biochemical stability, with no significant changes in any biomarkers during the *induction* phase. Although TBE did not induce statistically significant alterations, our data showed consistent trend toward reduction in both CK-MB and LDH.

Both supporting and divergent reports can be found in the literature (Ahiskalioglu et al., 2015; Ahiskalioglu et al., 2018; Chavda and Patel, 2022; Feng et al., 2013; Gil et al., 2004; Poon et al., 2000; Sanders et al., 1982; Trulson and Ulissey, 1987; Zorniak et al., 2010). Previous toxicology work showed that repeated oral exposure to CH produced time- and exposure-dependent changes in LDH activity, with both suppression and elevation in rodents (Poon et al., 2000; Sanders et al., 1982). In contrast, a single exposure protocol revealed no LDH changes in brains of rats (Trulson & Ulissey, 1987).

TP exhibited the least interference with cardiac biomarkers in our study. Ahiskalioglu et al. (2018) reported CK-MB values in rats that were comparable to our findings, with no elevation in LDH. Similarly, Gil et al. (2004) found unchanged plasma LDH levels following intravenous TP administration in rabbits. Zorniak et al. (2010) further demonstrated that TP induced the smallest increases in CK and LDH when compared with urethane and pentobarbital in rats.

In our study, KX was associated with an increase in CK-MB. Similarly, in diabetic rats subjected to middle cerebral artery occlusion to induce ischemic stroke, KX elevated serum CK-MB and LDH levels compared to controls. Interestingly, treatment with antidiabetic drugs, such as voglibose and saxagliptin reduced both biomarkers (Chavda and Patel, 2022).

Ketamine exposure has also been shown to increase cardiac CK-MB and cTnI levels in rat heart tissue. This effect was prevented by metyrosine, while metoprolol was only partially effective, suggesting that modulation of sympathetic activity can attenuate ketamine-induced on biomarkers elevation (Ahiskalioglu et al., 2015).

In contrast, TBE in our cohort did not induce statistically significant changes in LDH or cTnI. However, in a scald injury model, Feng et al. (2013) reported that TBE increased serum cTnI compared to sham animals, although the effect was less pronounced than that was observed with KX.

For ISO, our results showed no significant changes in cTnI, which aligns with clinical findings. Fellahi et al. (2004) reported that ISO did not alter cTnI release in patients undergoing cardiac surgery compared with the *pre-anesthetic* phase. Additional human studies have demonstrated cardioprotective effects of volatile anesthetics, including reductions in CK, CK-MB, and troponin-T following aortic cross-clamping and during early postoperative period (Ceyhan et al., 2011). These protective effects have been attributed to several mechanisms, including activation of protein kinase C epsilon, generation of reactive oxygen species (ROS), and opening of ATP-sensitive potassium channels (Lang et al., 2013; Oldenburg et al., 2002; Van Allen et al., 2012). Complementary experiments further support these cardioprotective properties. Liu et al. (2019) showed that sevoflurane reduced infarct size, LDH, and cTnI by mitigating endoplasmic reticulum stress via the PERK/eIF2α/ATF4 signaling pathway. Gong et al. (2012) further demonstrated that sevoflurane postconditioning decreased cTnI release, ROS production, and arrhythmia incidence in isolated rat hearts. More recently, Xu et al. (2024) reported that emulsified ISO pretreatment attenuated myocardial ischemia-reperfusion injury in rats, lowering CK-MB and cTnI levels and suppressing TLR-4 expression. Collectively, these studies indicate that both ISO and sevoflurane exert cardioprotective effects, reducing biomarkers of cardiac injury across experimental and clinical settings.

### 4.5. Limitations

A key limitation of this study is that enzymatic and blood-gas analyses were performed only during the *induction* phase, without additional measurements during recovery. This approach may have overlooked delayed or cumulative effects of the anesthetic agents on both cardiac biomarkers and respiratory parameters. Moreover, while conventional hemodynamic and biochemical endpoints were assessed, complementary molecular or histological analyses would be necessary to clarify the mechanisms underlying the observed alterations. Further studies should be necessary to investigate the long-term implications of our findings.

## 5. Conclusion

Our findings show that commonly used anesthetic agents exert heterogeneous effects on cardiorespiratory function and cardiac biomarkers. These results reinforce the need to select anesthetics based on their specific physiological impact, both to ensure methodological consistency in experimental studies and to avoid confounding effects on outcome measures. Importantly, the ability of some agents to modulate cardiac biomarkers also raises the possibility of cardioprotective properties, offering translational relevance for perioperative management in patients with cardiovascular risk.

## Supporting information

Statistical Summary Tables

## Data Availability Statement

All supporting data are available from the corresponding author upon request.

## Authors’ contributions

L.J.C. and V.S.M. performed biochemical assays and prepared the original draft of the manuscript. B.T.J. and B.A.A.M. carried out the in vivo assays. T.J.B. conducted biochemical assays together with L.J.C. and V.S.M. L.S. was responsible for variability data analysis, data interpretation, and validation. V.B. and D.A.M.G.B. were involved in the conceptualization of the study, data analysis, and funding acquisition. I.S.A.F. was involved in the conceptualization of the study, performed variability data analysis, contributed to manuscript review and editing and funding acquisition. K.N.S. was the lead responsible for the conceptualization of the study, and also carried out data analysis, supervision, funding acquisition, and manuscript review and editing. J.B.C, J.F.R.P. and F.D.M. contributed to data interpretation, provided resources, and participated in funding acquisition. All authors contributed to the critical review and editing of the manuscript, approved the final version, and agree to be accountable for all aspects of the work.

## References

Abdi-Azar, H., & Maleki, S. A. (2014). Comparison of the anesthesia with thiopental sodium alone and their combination with Citrus aurantium L. (Rutaseae) essential oil in male rat. Bulletin of Environment, Pharmacology and Life Sciences, 3(Special Issue V), 37–44.

Ahiskalioglu, A., Ince, I., Aksoy, M., Ahiskalioglu, E. O., Comez, M., Dostbil, A., Celik, M., Alp, H. H., Coskun, R., Taghizadehghalehjoughi, A., & Suleyman, B. (2015). Comparative Investigation of Protective Effects of Metyrosine and Metoprolol Against Ketamine Cardiotoxicity in Rats. Cardiovascular Toxicology, 15(4), 336–344. 10.1007/s12012-014-9301-z

Ahiskalioglu, E. O., Aydin, P., Ahiskalioglu, A., Suleyman, B., Kuyrukluyildiz, U., Kurt, N., Altuner, D., Coskun, R., & Suleyman, H. (2018). The effects of ketamine and thiopental used alone or in combination on the brain, heart, and bronchial tissues of rats. Archives of Medical Science, 14(3), 645–654. 10.5114/aoms.2016.59508

Ajadi, R. A., Gazal, N. A., Teketay, D. H., & Gazal, S. O. (2013). Evaluation of tribromoethanol, tribromoethanol-buprenorphine and ketamine-xylazine combinations for anaesthesia in sprague-dawley rats undergoing ovariectomy. Nigerian Journal of Physiological Sciences, 28(1), 51–56.

Albrecht, M., Henke, J., Tacke, S., Markert, M., & Guth, B. (2014). Effects of isoflurane, ketamine-xylazine and a combination of medetomidine, midazolam and fentanyl on physiological variables continuously measured by telemetry in Wistar rats. BMC Veterinary Research, 10(1), 1–14. 10.1186/s12917-014-0198-3

Aono, H., Hirakawa, M., Unruh, G. K., Kindscher, J. D., & Goto, H. (2001). Anesthetic induction agents, sympathetic nerve activity and baroreflex sensitivity: A study in rabbits comparing thiopental, propofol and etomidate. In Acta Medica Okayama (Vol. 55, Number 4, pp. 197–203). 10.18926/AMO/31994

Batista, T. J., Minassa, V. S., Aitken, A. V., Jara, B. T., Felippe, I. S. A., Beijamini, V., Paton, J. F. R., dos Santos, L., & Sampaio, K. N. (2019). Intermittent Exposure to Chlorpyrifos Differentially Impacts Neuroreflex Control of Cardiorespiratory Function in Rats. Cardiovascular Toxicology, 19(6), 548–564. 10.1007/s12012-019-09528-7

Brown, E. N., Purdon, P. L., & Van Dort, C. J. (2011). General anesthesia and altered states of arousal: A systems neuroscience analysis. Annual Review of Neuroscience, 34, 601–628. 10.1146/annurev-neuro-060909-153200

Brzezinski-Sinai, Y., Zwang, E., Plotnikova, E., Halizov, E., Shapira, I., Zeltser, D., Rogowski, O., Berliner, S., Matot, I., & Shenhar-Tsarfaty, S. (2021). Cholinesterase activity in serum during general anesthesia in patients with or without vascular disease. Scientific Reports, 11(1), 1–8. 10.1038/s41598-021-96251-5

Carrasco, M. S., Borbolla, M. G., Grájera, Á., Lozano, G. S., & Rahola, J. G. (1978). Influence of different anesthesics on the activity of serum pseudocholinesterase. Revista Española de Anestesiología y Reanimación, 25(2), 117–123.

Ceyhan, D., Tanriverdi, B., & Bilir, A. (2011). Comparison of the effects of sevoflurane and isoflurane on myocardial protection in coronary bypass surgery. Anadolu Kardiyoloji Dergisi/The Anatolian Journal of Cardiology, 257–263. 10.5152/akd.2011.059

Chavda, V., & Patel, S. (2022). Voglibose and saxagliptin ameliorate the post-surgical stress and cognitive dysfunction in chronic anaesthesia exposed diabetic MCAo induced ischemic rats. IBRO Neuroscience Reports, 13(November), 426–435. 10.1016/j.ibneur.2022.10.009

Costas-Ferreira, C., Durán, R., & Faro, L. F. (2023). Neurotoxic effects of exposure to glyphosate in rat striatum: Effects and mechanisms of action on dopaminergic neurotransmission. Pesticide Biochemistry and Physiology, 193(April). 10.1016/j.pestbp.2023.105433

Danese, E., & Montagnana, M. (2016). An historical approach to the diagnostic biomarkers of acute coronary syndrome. Annals of Translational Medicine, 4(10), 194–194. 10.21037/atm.2016.05.19

Deckardt, K., Weber, I., Kaspers, U., Hellwig, J., Tennekes, H., & van Ravenzwaay, B. (2007). The effects of inhalation anaesthetics on common clinical pathology parameters in laboratory rats. Food and Chemical Toxicology, 45(9), 1709–1718. 10.1016/j.fct.2007.03.005

Drorbaugh, J. E., & Fenn, W. O. (1955). A barometric method for measuring ventilation in newborn infants. Pediatrics, 16(1), 81–87. http://www.ncbi.nlm.nih.gov/pubmed/14394741

Eger, E. I. (2004). Characteristics of anesthetic agents used for induction and maintenance of general anesthesia. American Journal of Health-System Pharmacy, 61(SUPPL. 4), 3–10. 10.1093/ajhp/61.suppl_4.s3

Fazan, R., de Oliveira, M., Oliveira, J. A. C., Salgado, H. C., & Garcia-Cairasco, N. (2011). Changes in autonomic control of the cardiovascular system in the Wistar audiogenic rat (WAR) strain. Epilepsy & Behavior, 22(4), 666–670. 10.1016/j.yebeh.2011.09.010

Felippe, I. S. A., Müller, C. J. T., Siqueira, A. A., dos Santos, L., Cavadino, A., Paton, J. F. R., Beijamini, V., & Sampaio, K. N. (2020). The antidotes atropine and pralidoxime distinctively recover cardiorespiratory components impaired by acute poisoning with chlorpyrifos in rats. Toxicology and Applied Pharmacology, 389(October 2019), 114879. 10.1016/j.taap.2020.114879

Fellahi, J. L., Gue, X., Philippe, E., Riou, B., & Gerard, J. L. (2004). Isoflurane may noy influence postoperative cardiac troponin I release and clinical outcome in adult cardiac surgery. European Journal of Anaesthesiology, 21, 688–693.

Feng, Y., Chai, J., Chu, W., Ma, L., Zhang, P., & Duan, H. (2013). Combination of ketamine and xylazine exacerbates cardiac dysfunction in severely scalded rats during the shock stage. Experimental and Therapeutic Medicine, 6(3), 641–648. 10.3892/etm.2013.1213

Field, K. J., White, W. J., & Lang, C. M. (1993). Anaesthetic effects of chloral hydrate, pentobarbitone and urethane in adult male rats. Laboratory Animals, 27(3), 258–269. 10.1258/002367793780745471

Flecknell, P. (2016). Laboratory Animal Anaesthesia (4th ed.). Academic Press. 10.1016/B978-0-12-800036-6.00001-6

Flecknell, P. A. (2009a). Anaesthesia of Common Laboratory Species: Special Considerations. In Laboratory Animal Anaesthesia (pp. 181–241). 10.1016/b978-0-12-369376-1.00006-x

Flecknell, P. A. (2009b). Anaesthetic Management. In Laboratory Animal Anaesthesia (3rd ed., pp. 79–108). 10.1016/B978-0-12-369376-1.00003-4

Franks, N. P. (2008). General anaesthesia: From molecular targets to neuronal pathways of sleep and arousal. Nature Reviews Neuroscience, 9(5), 370–386. 10.1038/nrn2372

Garg, P., Morris, P., Fazlanie, A. L., Vijayan, S., Dancso, B., Dastidar, A. G., Plein, S., Mueller, C., & Haaf, P. (2017). Cardiac biomarkers of acute coronary syndrome: from history to high-sensitivity cardiac troponin. Internal and Emergency Medicine, 12(2), 147–155. 10.1007/s11739-017-1612-1

Gargiulo, S., Albanese, S., Megna, R., Gramanzini, M., Marsella, G., & Vecchiarelli, L. (2025). Veterinary medical care in rodent models of stroke: Pitfalls and refinements to balance quality of science and animal welfare. Neuroscience, 572(September 2024), 269–302. 10.1016/j.neuroscience.2025.01.044

Gargiulo, S., Greco, A., Gramanzini, M., Esposito, S., Affuso, A., Brunetti, A., & Vesce, G. (2012). Mice Anesthesia, Analgesia, and Care, Part I: Anesthetic Considerations in Preclinical Research. ILAR Journal, 53(1), E55–E69. 10.1093/ilar.53.1.55

Gil, A. G., Silván, G., Illera, M., & Illera, J. C. (2004). The effects of anesthesia on the clinical chemistry of New Zealand White rabbits. Contemporary Topics in Laboratory Animal Science, 43(3), 25–29.

Goldstein, A., & Aronow, L. (1959). The durations of action of thiopental and pentobarbital. 128, 6.

Goliasch, G., Haschemi, A., Marculescu, R., Endler, G., Maurer, G., Wagner, O., Huber, K., Mannhalter, C., & Niessner, A. (2012). Butyrylcholinesterase Activity Predicts Long-Term Survival in Patients with Coronary Artery Disease. Clinical Chemistry, 58(6), 1055–1058. 10.1373/clinchem.2011.175984

Gonca, E. (2015). Comparison of thiopental and ketamine+xylazine anesthesia in ischemia/reperfusion-induced arrhythmias in rats. Turkish Journal of Medical Sciences, 45(6), 1413–1420. 10.3906/sag-1403-25

Gong, J.-S., Yao, Y.-T., Fang, N.-X., & Li, L.-H. (2012). Sevoflurane postconditioning attenuates reperfusion-induced ventricular arrhythmias in isolated rat hearts exposed to ischemia/reperfusion injury. Molecular Biology Reports, 39(6), 6417–6425. 10.1007/s11033-012-1447-9

Greene, S. A., & Thurmon, J. C. (1988). Xylazine – a review of its pharmacology and use in veterinary medicine. Journal of Veterinary Pharmacology and Therapeutics, 11(4), 295–313. 10.1111/j.1365-2885.1988.tb00189.x

Guyenet, P. G., Stornetta, R. L., Holloway, B. B., Souza, G. M. P. R., & Abbott, S. B. G. (2018). Rostral Ventrolateral Medulla and Hypertension. Hypertension, 72(3), 559–566. 10.1161/HYPERTENSIONAHA.118.10921

Haskins, S. C., Farver, T. B., & Patz, J. D. (1985). Ketamine in dogs. Am J Vet Res, 46(9), 1855–1860.

Heistein, L. C. (2006). Chloral Hydrate Sedation for Pediatric Echocardiography: Physiologic Responses, Adverse Events, and Risk Factors. PEDIATRICS, 117(3), e434–e441. 10.1542/peds.2005-1445

Hemmings, H. C., Riegelhaupt, P. M., Kelz, M. B., Solt, K., Eckenhoff, R. G., Orser, B. A., & Goldstein, P. A. (2019). Towards a Comprehensive Understanding of Anesthetic Mechanisms of Action: A Decade of Discovery. Trends in Pharmacological Sciences, 40(7), 464–481. 10.1016/j.tips.2019.05.001

Herbst, L. S., Gaigher, T., Siqueira, A. A., Joca, S. R. L., Sampaio, K. N., & Beijamini, V. (2019). New evidence for refinement of anesthetic choice in procedures preceding the forced swimming test and the elevated plus-maze. Behavioural Brain Research, 368(April), 111897. 10.1016/j.bbr.2019.04.011

Hørder, M., Magid, E., Pitkänen, E., Härkönen, M., Strömme, J. H., Theodorsen, L., Gerhardt, W., & Waldenström, J. (1979). Recommended method for the determination of creatine kinase in blood modified by the inclusion of EDTA: The Committee on Enzymes of The Scandinavian Society for Clinical Chemistry and Clinical Physiology (SCE). Scandinavian Journal of Clinical and Laboratory Investigation, 39(1), 1–5. 10.3109/00365517909104932

Keshavarz, S., Nemati, M., Saied Salehi, M., & Naseh, M. (2023). The impact of anesthetic drugs on hemodynamic parameters and neurological outcomes following temporal middle cerebral artery occlusion in rats. NeuroReport, 34(4), 199–204. 10.1097/WNR.0000000000001863

Khan, K. S., Hayes, I., & Buggy, D. J. (2014). Pharmacology of anaesthetic agents I: Intravenous anaesthetic agents. Continuing Education in Anaesthesia, Critical Care and Pain, 14(3), 100–105. 10.1093/bjaceaccp/mkt039

Kushawaha, S., Malpani, A., & Aswar, U. (2011). Effect of different anaesthetic agents on cardiovascular parameters in male Wistar rats. Research Journal of Pharmaceutical, Biological and Chemical Sciences, 2(2), 685–690.

Lang, X. E., Wang, X., Zhang, K. R., Lv, J. Y., Jin, J. H., & Li, Q. S. (2013). Isoflurane Preconditioning Confers Cardioprotection by Activation of ALDH2. PLoS ONE, 8(2). 10.1371/journal.pone.0052469

Létienne, R., Bel, L., Bessac, A. M., Denais, D., Degryse, A. D., John, G. W., & Le Grand, B. (2006). Cardioprotection of cariporide evaluated by plasma myoglobin and troponin I in myocardial infarction in pigs. Fundamental and Clinical Pharmacology, 20(2), 105–113. 10.1111/j.1472-8206.2006.00394.x

Lieggi, C. C., Fortman, J. D., Kleps, R. A., Sethi, V., Anderson, J. A., Brown, C. E., & Artwohl, J. E. (2005). An evaluation of preparation methods and storage conditions of tribromoethanol. Contemporary Topics in Laboratory Animal Science, 44(1), 11–16. http://www.ncbi.nlm.nih.gov/pubmed/15697192

Liu, A.-J., Pang, C.-X., Liu, G.-Q., Wang, S.-D., Chu, C.-Q., Li, L.-Z., Dong, Y., & Zhu, D.-Z. (2019). Ameliorative effect of sevoflurane on endoplasmic reticulum stress mediates cardioprotection against ischemia–reperfusion injury. Canadian Journal of Physiology and Pharmacology, 97(5), 345–351. 10.1139/cjpp-2018-0016

Massey, C. A., Iceman, K. E., Johansen, S. L., Wu, Y., Harris, M. B., & Richerson, G. B. (2015). Isoflurane abolishes spontaneous firing of serotonin neurons and masks their pH/CO2 chemosensitivity. Journal of Neurophysiology, 113(7), 2879–2888. 10.1152/jn.01073.2014

Massey, C. A., & Richerson, G. B. (2017). Isoflurane, ketamine-xylazine, and urethane markedly alter breathing even at subtherapeutic doses. Journal of Neurophysiology, 118(4), 2389–2401. 10.1152/jn.00350.2017

Maud, P., Thavarak, O., Cédrick, L., Michèle, B., Vincent, B., Olivier, P., & Régis, B. (2014). Evidence for the Use of Isoflurane as a Replacement for Chloral Hydrate Anesthesia in Experimental Stroke: An Ethical Issue. BioMed Research International, 2014, 1–7. 10.1155/2014/802539

Meyer, R. E., & Fish, R. E. (2005a). A review of tribromoethanol anesthesia for production of genetically engineered mice and rats. Lab Animal, 34(10), 47–52. 10.1038/laban1105-47

Meyer, R. E., & Fish, R. E. (2005b). A review of tribromoethanol anesthesia for production of genetically engineered mice and rats. In Lab Animal (Vol. 34, Number 10, pp. 47–52). 10.1038/laban1105-47

Michels, B., Holzamer, A., Graf, B. M., Bredthauer, A., Petermichl, W., Müller, A., Zausig, Y. A., & Bitzinger, D. I. (2021). Butyrylcholinesterase as a perioperative complication marker in patients after transcatheter aortic valve implantation: A prospective observational study. BMJ Open, 11(7), 1–9. 10.1136/bmjopen-2020-042857

Morton, L. D., Sanders, M., Reagan, W. J., Crabbs, T. A., McVean, M., & Funk, K. A. (2019). Confounding Factors in the Interpretation of Preclinical Studies. International Journal of Toxicology, 38(3), 228–234. 10.1177/1091581819837157

Mosneag, I. E., Flaherty, S. M., Wykes, R. C., & Allan, S. M. (2024). Stroke and Translational Research – Review of Experimental Models with a Focus on Awake Ischaemic Induction and Anaesthesia. Neuroscience, 550(November 2023), 89–101. 10.1016/j.neuroscience.2023.11.034

Nana, A., Cardan, E., & Cucuianu, M. (1977). Pseudocholinesterase changes in anesthesia using pancuronium. Acta Anaesthesiologica Belgica, 28(3), 183–187. http://www.ncbi.nlm.nih.gov/pubmed/612115

Navarro, K. L., Huss, M., Smith, J. C., Sharp, P., Marx, J. O., & Pacharinsak, C. (2021). Mouse Anesthesia: The Art and Science. ILAR Journal, 62(1–2), 238–273. 10.1093/ilar/ilab016

Ochiai, Y., Iwano, H., Sakamoto, T., Hirabayashi, M., Kaneko, E., Watanabe, T., Yamashita, K., & Yokota, H. (2016). Blood biochemical changes in mice after administration of a mixture of three anesthetic agents. Journal of Veterinary Medical Science, 78(6), 951–956. 10.1292/jvms.15-0474

Oh, S. S., & Narver, H. L. (2024). Mouse and Rat Anesthesia and Analgesia. Current Protocols, 4(2). 10.1002/cpz1.995

Oldenburg, O., Cohen, M. V., Yellon, D. M., & Downey, J. M. (2002). Mitochondrial KATP channels: Role in cardioprotection. In Cardiovascular Research (Vol. 55, Number 3, pp. 429–437). 10.1016/S0008-6363(02)00439-X

Olmos-Pastoresa, C. A., Vázquez-Mendoza, E., López-Meraz, M. L., Pérez-Estudillo, C. A., Beltran-Parrazal, L., & Morgado-Valle, C. (2023). Transgenic rodents as dynamic models for the study of respiratory rhythm generation and modulation: a scoping review and a bibliometric analysis. Frontiers in Physiology, 14(December), 1–25. 10.3389/fphys.2023.1295632

Pachon, R. E., Scharf, B. A., Vatner, D. E., & Vatner, S. F. (2015). Best anesthetics for assessing left ventricular systolic function by echocardiography in mice. American Journal of Physiology-Heart and Circulatory Physiology, 308(12), H1525–H1529. 10.1152/ajpheart.00890.2014

Parikh, M., & Pierce, G. N. (2024). Considerations for choosing an optimal animal model of cardiovascular disease. Canadian Journal of Physiology and Pharmacology, 102(2), 75–85. 10.1139/cjpp-2023-0206

Poon, R., Nadeau, B., & Chu, I. (2000). Biochemical effects of chloral hydrate on male rats following 7-day drinking water exposure. Journal of Applied Toxicology, 20(6), 455–461. 10.1002/1099-1263(200011/12)20:6<455::AID-JAT714>3.0.CO;2-A

Sanders, V. M., Kauffmann, B. M., & White, K. L. (1982). Toxicology of chloral hydrate in the mouse. Environmental Health Perspectives, Vol. 44(April), 137–146. 10.1289/ehp.8244137

Saraswat, V. (2015). Effects of anaesthesia techniques and drugs on pulmonary function. Indian Journal of Anaesthesia, 59(9), 557–564. 10.4103/0019-5049.165850

Schmidt, et al. (1992). Proposal of Standard Methods for Determination of Enzymatic Catalytic Concentrations in Serum and Plasma at 37°C. Eur. J. Clin. Chem. Clin. Biochem., 30(3), 163–170.

Schumann, G., Bonora, R., Ceriotti, F., Clerc-Renaud, P., Ferrero, C. A., Férard, G., Franck, P. F. H., Gella, F.-J., Hoelzel, W., Jørgensen, P. J., Kanno, T., Kessner, A., Klauke, R., Kristiansen, N., Lessinger, J.-M., Linsinger, T. P. J., Misaki, H., & Panteghini, M. (2002). IFCC Primary Reference Procedures for the Measurement of Catalytic Activity Concentrations of Enzymes at 37°C. Part 3. Reference Procedure for the Measurement of Catalytic Concentration of Lactate Dehydrogenase. Clinical Chemistry and Laboratory Medicine, 40(6). 10.1515/CCLM.2002.111

Schwarzkopf, T. M., Horn, T., Lang, D., & Klein, J. (2013). Blood gases and energy metabolites in mouse blood before and after cerebral ischemia: The effects of anesthetics. Experimental Biology and Medicine, 238(1), 84–89. 10.1258/ebm.2012.012261

Shcherbak, N. S., Yukina, G. Y., Gurbo, A. G., Sukhorukova, E. G., Sargsian, A. G., & Tomson, V. V. (2024). Effects of Chloral Hydrate on the Morphogenetic Characteristics of the Neocortex and Functional Status in Elderly Male Rats. Neuroscience and Behavioral Physiology, 54(3), 474–481. 10.1007/s11055-024-01614-w

Shrestha, J., Paudel, K. R., Nazari, H., Dharwal, V., Bazaz, S. R., Johansen, M. D., Dua, K., Hansbro, P. M., & Warkiani, M. E. (2023). Advanced models for respiratory disease and drug studies. Medicinal Research Reviews, 43(5), 1470–1503. 10.1002/med.21956

Sinai, Y. B., Zwang, E., Plotnikova, E., Halizov, E., Shapira, I., Zeltser, D., Rogowski, O., Berliner, S., Matot, I., & Tsarfaty, S. S. (2021). Cholinesterase activity in serum during general anesthesia in patients with or without vascular disease. Scientific Reports, 11, 1–8.

Sisson, D. F., & Siegel, J. (1989). Chloral hydrate anesthesia: EEG power spectrum analysis and effects on VEPs in the rat. Neurotoxicology and Teratology, 11(1), 51–56. 10.1016/0892-0362(89)90085-8

Sumitra, M., Manikandan, P., Rao, K. V. K., Nayeem, M., Manohar, B. M., & Puvanakrishnan, R. (2004). Cardiorespiratory effects of diazepam-ketamine, xylazine-ketamine and thiopentone anesthesia in male Wistar rats - A comparative analysis. Life Sciences, 75(15), 1887–1896. 10.1016/j.lfs.2004.05.009

Sun, L., Qi, X., Tan, Q., Yang, H., & Qi, X. (2016). Low Serum-Butyrylcholinesterase Activity as a Prognostic Marker of Mortality Associates with Poor Cardiac Function in Acute Myocardial Infarction. Clinical Laboratory, 62(6), 1093–1099. http://www.ncbi.nlm.nih.gov/pubmed/27468571

Thygesen, K., Alpert, J. S., & White, H. D. (2007). Universal Definition of Myocardial Infarction. Circulation, 116(22), 2634–2653. 10.1161/CIRCULATIONAHA.107.187397

Trulson, M. E., & Ulissey, M. J. (1987). Chloral hydrate anesthesia alters cerebral enzymes in the rat. A histochemical study. Acta Anatomica, 130(4), 319–323. http://www.ncbi.nlm.nih.gov/pubmed/3434184

Vachon, P., Faubert, S., Blais, D., Comtois, A., & Bienvenu, J. G. (2000). A pathophysiological study of abdominal organs following intraperitoneal injections of chloral hydrate in rats: comparison between two anaesthesia protocols. Laboratory Animals, 34(1), 84–90. 10.1258/002367700780578082

Van Allen, N. R., Krafft, P. R., Leitzke, A. S., Applegate, R. L., Tang, J., & Zhang, J. H. (2012). The role of Volatile Anesthetics in Cardioprotection: a systematic review. Medical Gas Research, 2(1), 22. 10.1186/2045-9912-2-22

van Doorn, E. C. H., Amesz, J. H., Sadeghi, A. H., de Groot, N. M. S., Manintveld, O. C., & Taverne, Y. J. H. J. (2024). Preclinical Models of Cardiac Disease: A Comprehensive Overview for Clinical Scientists. Cardiovascular Engineering and Technology, 15(2), 232–249. 10.1007/s13239-023-00707-w

Vardanyan, R. S., & Hruby, V. J. (2006). Synthesis of Essential Drugs. In Elsevier (1st ed.). Elsevier. 10.1055/s-0031-1296659

Wada, D. R., Harashima, H., Ebling, W., Osaki, E. W., & Stanski, D. R. (1996). Effects of thiopental on regional blood flows in the rat. Anesthesiology, 84(3), 596–604. http://www.ncbi.nlm.nih.gov/pubmed/8659788

Ward-Flanagan, R., & Dickson, C. T. (2023). Intravenous chloral hydrate anesthesia provides appropriate analgesia for surgical interventions in male Sprague-Dawley rats. PLoS ONE, 18(6 June), 1–10. 10.1371/journal.pone.0286504

Weiss, J., & Zimmermann, F. (1999). Tribromoethanol (Avertin) as an anaesthetic in mice. Laboratory Animals, 33(2), 192–193. 10.1258/002367799780578417

Wellington, D., Mikaelian, I., & Singer, L. (2013). Comparison of Ketamine – Xylazine and Ketamine – Dexmedetomidine Anesthesia and Intraperitoneal Tolerance in Rats. 52(4), 481–487.

Xu, Z., Li, Z., Chen, S., Zhu, Y., Wang, Y., Zhan, J., & Wu, Y. (2024). Emulsified isoflurane pretreatment attenuates myocardial ischemia-reperfusion injuries by suppressing toll-like Receptor-4. Immunopharmacology and Immunotoxicology, 46(6), 751–756. 10.1080/08923973.2024.2399266

Zatroch, K. K., Knight, C. G., Reimer, J. N., & Pang, D. S. J. (2016). Refinement of intraperitoneal injection of sodium pentobarbital for euthanasia in laboratory rats (Rattus norvegicus). BMC Veterinary Research, 13(1), 60. 10.1186/s12917-017-0982-y

Zausinger, S., Baethmann, A., & Schmid-Elsaesser, R. (2002). Anesthetic methods in rats determine outcome after experimental focal cerebral ischemia: Mechanical ventilation is required to obtain controlled experimental conditions. Brain Research Protocols, 9(2), 112–121. 10.1016/S1385-299X(02)00138-1

Zhang, X., Yi, Y., Cheng, L., Chen, H., & Hu, Y. (2023). Dynamic effects of miR-20a-5p on hippocampal ripple energy after status epilepticus in rats. Experimental Brain Research, 241(8), 2097–2106. 10.1007/s00221-023-06663-0

Zivkovic, A. R., Schmidt, K., Sigl, A., Decker, S. O., Brenner, T., & Hofer, S. (2015). Reduced Serum Butyrylcholinesterase Activity Indicates Severe Systemic Inflammation in Critically Ill Patients. 2015. 10.1155/2015/274607

Zorniak, M., Mitrega, K., Bialka, S., Porc, M., & Krzeminski, T. F. (2010). Comparison of thiopental, urethane, and pentobarbital in the study of experimental cardiology in rats in vivo. J Cardiovasc Pharmacol, 56(1), 38–44. 10.1097/FJC.0b013e3181dd502c

