## Supplementary material for "Cardiorespiratory and Cardiac Biomarker Responses to Five Anesthetic Regimens in Rats": Statistical Summary Tables

**ORCID:**

Vanessa Beijamini 0000-0003-4533-0495

Igor Simões Assunção Felippe 0000-0002-8582-2104

Karla N. Sampaio 0000-0003-0293-0482

*L.J.C., V.S.M and B.T.J. are equal first authors.*

^‡^**Corresponding authors:**

**Professor Karla Nívea Sampaio**: Federal University of Espírito Santo; Postgraduate Program in Pharmaceutical Sciences; Department of Pharmaceutical Sciences, Health Sciences Center; Av. Marechal Campos, 1468, Maruípe, Vitória, ES, Brazil; Postcode: 29047-105;; Phone Number: +55 27 99933-2446;

**Dr. Igor Simões Assunção Felippe**: The Centre for Heart Research – Manaaki Mānawa, Department of Physiology, Faculty of Health & Medical Sciences, University of Auckland, Grafton Campus, Auckland, 1023, New Zealand.; Phone number: +64 22 495 3593

**Appendix 1**

Statistical Summary Tables

**Table A1**. Supplementary Statistical Analyses and Results: Hemodynamic Data, Induction phase

| **Δ SBP (mmHg)** |  |  |  |
| --- | --- | --- | --- |
| Tukey's multiple comparisons test | Mean Differences | CI95% adjust | P Value |
| CH vs. KX | -72.09 | -91.65 to -52.52 | <0.0001 |
| CH vs. TBE | -26.03 | -45.17 to -6.901 | 0.003 |
| CH vs. TP | -30.26 | -50.33 to -10.20 | 0.0008 |
| CH vs. ISO | 10.91 | -8.649 to 30.48 | 0.5183 |
| KX vs. TBE | 46.05 | 26.49 to 65.62 | <0.0001 |
| KX vs. TP | 41.83 | 21.35 to 62.30 | <0.0001 |
| KX vs. ISO | 83.0 | 63.02 to 103.0 | <0.0001 |
| TBE vs. TP | -4.228 | -24.29 to 15.84 | 0.9751 |
| TBE vs. ISO | 36.95 | 17.38 to 56.51 | <0.0001 |
| TP vs. ISO | 41.18 | 20.70 to 61.65 | <0.0001 |
| **Δ DBP (mmHg)** |  |  |  |
| Tukey's multiple comparisons test | Mean Differences | CI95% adjust | P Value |
| CH vs. KX | -54.55 | -69.50 to -39.59 | <0.0001 |
| CH vs. TBE | -23.95 | -38.58 to -9.324 | 0.0002 |
| CH vs. TP | -29.65 | -44.99 to -14.30 | <0.0001 |
| CH vs. ISO | 8.35 | -6.606 to 23.31 | 0.5176 |
| KX vs. TBE | 30.6 | 15.64 to 45.55 | <0.0001 |
| KX vs. TP | 24.9 | 9.249 to 40.56 | 0.0004 |
| KX vs. ISO | 62.9 | 47.62 to 78.18 | <0.0001 |
| TBE vs. TP | -5.694 | -21.04 to 9.647 | 0.831 |
| TBE vs. ISO | 32.3 | 17.35 to 47.26 | <0.0001 |
| TP vs. ISO | 38.0 | 22.34 to 53.65 | <0.0001 |
| **Δ MAP (mmHg)** |  |  |  |
| Tukey's multiple comparisons test | Mean Differences | CI95% adjust | P Value |
| CH vs. KX | -61.19 | -77.46 to -44.93 | <0.0001 |
| CH vs. TBE | -24.5 | -40.40 to -8.588 | 0.0006 |
| CH vs. TP | -30.07 | -46.75 to -13.38 | <0.0001 |
| CH vs. ISO | 9.426 | -6.840 to 25.69 | 0.4803 |
| KX vs. TBE | 36.7 | 20.43 to 52.96 | <0.0001 |
| KX vs. TP | 31.13 | 14.10 to 48.15 | <0.0001 |
| KX vs. ISO | 70.62 | 54.00 to 87.24 | <0.0001 |
| TBE vs. TP | -5.57 | -22.25 to 11.11 | 0.8782 |
| TBE vs. ISO | 33.92 | 17.66 to 50.19 | <0.0001 |
| TP vs. ISO | 39.49 | 22.47 to 56.52 | <0.0001 |
| **Δ HR (bpm)** |  |  |  |
| Tukey's multiple comparisons test | Mean Differences | CI95% adjust | P Value |
| CH vs. KX | 112.6 | 70.53 to 154.6 | <0.0001 |
| CH vs. TBE | -12.79 | -53.91 to 28.33 | 0.9032 |
| CH vs. TP | 22.41 | -20.72 to 65.53 | 0.5867 |
| CH vs. ISO | 44.98 | 2.933 to 87.02 | 0.0304 |
| KX vs. TBE | -125.4 | -167.4 to -83.31 | <0.0001 |
| KX vs. TP | -90.17 | -134.2 to -46.16 | <0.0001 |
| KX vs. ISO | -67.6 | -110.5 to -24.65 | 0.0004 |
| TBE vs. TP | 35.19 | -7.934 to 78.32 | 0.1592 |
| TBE vs. ISO | 57.76 | 15.72 to 99.81 | 0.0026 |
| TP vs. ISO | 22.57 | -21.44 to 66.58 | 0.5988 |

Mean difference = left group − right group. Positive values indicate a higher mean in the first group; negative values indicate a higher mean in the second group.

**Table A2**. Supplementary Statistical Analyses and Results: Hemodynamic Data, Recovery phase

| **Δ SBP (mmHg)** |  |  |  |
| --- | --- | --- | --- |
| Tukey's multiple comparisons test | Mean Differences | CI95% adjust | P Value |
| CH vs. KX | -39.18 | -55.58 to -22.78 | <0.0001 |
| CH vs. TBE | -37.45 | -53.49 to -21.41 | <0.0001 |
| CH vs. TP | -35.98 | -52.80 to -19.15 | <0.0001 |
| CH vs. ISO | -22.27 | -38.67 to -5.865 | 0.003 |
| KX vs. TBE | 1.736 | -14.66 to 18.14 | 0.9982 |
| KX vs. TP | 3.209 | -13.96 to 20.38 | 0.984 |
| KX vs. ISO | 16.92 | 0.1667 to 33.67 | 0.0467 |
| TBE vs. TP | 1.473 | -15.35 to 18.30 | 0.9991 |
| TBE vs. ISO | 15.18 | -1.216 to 31.58 | 0.082 |
| TP vs. ISO | 13.71 | -3.456 to 30.88 | 0.1754 |
| **Δ DBP (mmHg)** |  |  |  |
| Tukey's multiple comparisons test | Mean Differences | CI95% adjust | P Value |
| CH vs. KX | -30.74 | -45.56 to -15.93 | <0.0001 |
| CH vs. TBE | -32.95 | -47.43 to -18.46 | <0.0001 |
| CH vs. TP | -31.74 | -47.94 to -15.55 | <0.0001 |
| CH vs. ISO | -23.23 | -38.04 to -8.412 | 0.0005 |
| KX vs. TBE | -2.203 | -17.02 to 12.61 | 0.9932 |
| KX vs. TP | -1.001 | -17.49 to 15.49 | 0.9998 |
| KX vs. ISO | 7.518 | -7.614 to 22.65 | 0.6262 |
| TBE vs. TP | 1.202 | -15.00 to 17.40 | 0.9996 |
| TBE vs. ISO | 9.721 | -5.092 to 24.53 | 0.353 |
| TP vs. ISO | 8.519 | -7.970 to 25.01 | 0.5908 |
| **Δ MAP (mmHg)** |  |  |  |
| Tukey's multiple comparisons test | Mean Differences | CI95% adjust | P Value |
| CH vs. KX | -34.72 | -50.15 to -19.29 | <0.0001 |
| CH vs. TBE | -35.09 | -50.18 to -20.00 | <0.0001 |
| CH vs. TP | -34.11 | -49.94 to -18.28 | <0.0001 |
| CH vs. ISO | -22.73 | -38.16 to -7.301 | 0.0011 |
| KX vs. TBE | -0.3683 | -15.80 to 15.06 | >0.9999 |
| KX vs. TP | 0.6159 | -15.54 to 16.77 | >0.9999 |
| KX vs. ISO | 11.99 | -3.770 to 27.75 | 0.2149 |
| TBE vs. TP | 0.9842 | -14.84 to 16.81 | 0.9998 |
| TBE vs. ISO | 12.36 | -3.070 to 27.79 | 0.1731 |
| TP vs. ISO | 11.38 | -4.775 to 27.53 | 0.2846 |
| **Δ HR (bpm)** |  |  |  |
| Tukey's multiple comparisons test | Mean Differences | CI95% adjust | P Value |
| CH vs. KX | 57.34 | -6.857 to 121.5 | 0.1008 |
| CH vs. TBE | -74.61 | -137.4 to -11.83 | 0.0123 |
| CH vs. TP | -31.5 | -97.35 to 34.34 | 0.6599 |
| CH vs. ISO | -35.46 | -99.66 to 28.73 | 0.528 |
| KX vs. TBE | -131.9 | -196.1 to -67.75 | <0.0001 |
| KX vs. TP | -88.84 | -156.0 to -21.65 | 0.0041 |
| KX vs. ISO | -92.8 | -158.4 to -27.22 | 0.0018 |
| TBE vs. TP | 43.11 | -22.74 to 109.0 | 0.3565 |
| TBE vs. ISO | 39.15 | -25.05 to 103.3 | 0.4285 |
| TP vs. ISO | -3.958 | -71.15 to 63.24 | 0.9998 |

Mean difference = left group − right group. Positive values indicate a higher mean in the first group; negative values indicate a higher mean in the second group.

**Table A3.** Supplementary Statistical Analyses and Results: Respiratory Data, Induction phase

| **Δ VE (mL.Kg^-1^.min^-1^)** |  |  |  |
| --- | --- | --- | --- |
| Tukey's multiple comparisons test | Mean Differences | CI95% adjust | P Value |
| CH vs. KX | 54.01 | -58.57 to 166.6 | 0.5785 |
| CH vs. TBE | 4.572 | -105.9 to 115.1 | 0.9995 |
| CH vs. TP | -29.96 | -147.8 to 87.89 | 0.9041 |
| KX vs. TBE | -49.44 | -157.4 to 58.53 | 0.6148 |
| KX vs. TP | -83.97 | -199.5 to 31.51 | 0.2253 |
| TBE vs. TP | -34.53 | -148.0 to 78.92 | 0.8474 |
| **Δ *f*_R_ (cpm)** |  |  |  |
| Tukey's multiple comparisons test | Mean Differences | CI95% adjust | P Value |
| CH vs. KX | 8.541 | -17.98 to 35.06 | 0.8245 |
| CH vs. TBE | -20.48 | -46.50 to 5.549 | 0.1683 |
| CH vs. TP | -9.109 | -36.87 to 18.65 | 0.8163 |
| KX vs. TBE | -29.02 | -54.45 to -3.587 | 0.0198 |
| KX vs. TP | -17.65 | -44.85 to 9.551 | 0.3185 |
| TBE vs. TP | 11.37 | -15.35 to 38.09 | 0.6684 |
| **Δ VT (mL.Kg^-1^)** |  |  |  |
| Tukey's multiple comparisons test | Mean Differences | CI95% adjust | P Value |
| CH vs. KX | 0.1932 | -0.4869 to 0.8733 | 0.8719 |
| CH vs. TBE | 0.592 | -0.07541 to 1.259 | 0.0982 |
| CH vs. TP | 0.2042 | -0.5077 to 0.9160 | 0.8687 |
| KX vs. TBE | 0.3988 | -0.2534 to 1.051 | 0.3701 |
| KX vs. TP | 0.011 | -0.6866 to 0.7086 | >0.9999 |
| TBE vs. TP | -0.3878 | -1.073 to 0.2974 | 0.4384 |

Mean difference = left group − right group. Positive values indicate a higher mean in the first group; negative values indicate a higher mean in the second group.

**Table A4.** Supplementary Statistical Analyses and Results: Respiratory Data, Recovery phase

| **Δ VE (mL.Kg^-1^.min^-1^)** |  |  |  |
| --- | --- | --- | --- |
| Tukey's multiple comparisons test | Mean Differences | CI95% adjust | P Value |
| CH vs. KX | 25.17 | -99.10 to 149.4 | 0.9482 |
| CH vs. TBE | -24.81 | -146.8 to 97.15 | 0.9476 |
| CH vs. TP | -10.56 | -140.6 to 119.5 | 0.9963 |
| KX vs. TBE | -49.98 | -169.2 to 69.20 | 0.6783 |
| KX vs. TP | -35.73 | -163.2 to 91.74 | 0.8763 |
| TBE vs. TP | 14.25 | -111.0 to 139.5 | 0.9901 |
| **Δ *f*_R_ (cpm)** |  |  |  |
| Tukey's multiple comparisons test | Mean Differences | CI95% adjust | P Value |
| CH vs. KX | -2.046 | -30.30 to 26.21 | 0.9974 |
| CH vs. TBE | -10.47 | -38.21 to 17.26 | 0.7444 |
| CH vs. TP | 11.9 | -17.68 to 41.47 | 0.7059 |
| KX vs. TBE | -8.426 | -35.53 to 18.67 | 0.8391 |
| KX vs. TP | 13.94 | -15.04 to 42.93 | 0.5763 |
| TBE vs. TP | 22.37 | -6.105 to 50.84 | 0.1693 |
| **Δ VT (mL.Kg^-1^)** |  |  |  |
| Tukey's multiple comparisons test | Mean Differences | CI95% adjust | P Value |
| CH vs. KX | 0.4121 | -0.3224 to 1.147 | 0.446 |
| CH vs. TBE | -0.0207 | -0.7416 to 0.7002 | 0.9998 |
| CH vs. TP | -0.2785 | -1.047 to 0.4903 | 0.7676 |
| KX vs. TBE | -0.4328 | -1.137 to 0.2716 | 0.366 |
| KX vs. TP | -0.6907 | -1.444 to 0.06275 | 0.0828 |
| TBE vs. TP | -0.2578 | -0.9980 to 0.4823 | 0.788 |

Mean difference = left group − right group. Positive values indicate a higher mean in the first group; negative values indicate a higher mean in the second group.

**Table A5.** Supplementary Statistical Analyses and Results: Baroreflex Effectiveness Index, Induction and Recovery phases

| **Δ BEI: Induction** |  |  |  |
| --- | --- | --- | --- |
| Tukey's multiple comparisons test | Mean Differences | CI95% adjust | P Value |
| CH vs. KX | 0.12 | 0.007487 to 0.2326 | 0.0315 |
| CH vs. TBE | 0.05204 | -0.05163 to 0.1557 | 0.6119 |
| CH vs. TP | 0.06428 | -0.04826 to 0.1768 | 0.4887 |
| CH vs. ISO | -0.0229 | -0.1290 to 0.08320 | 0.9719 |
| KX vs. TBE | -0.06799 | -0.1782 to 0.04225 | 0.4113 |
| KX vs. TP | -0.05575 | -0.1744 to 0.06288 | 0.6687 |
| KX vs. ISO | -0.1429 | -0.2555 to -0.03039 | 0.0067 |
| TBE vs. TP | 0.01224 | -0.09800 to 0.1225 | 0.9977 |
| TBE vs. ISO | -0.07494 | -0.1786 to 0.02873 | 0.2564 |
| TP vs. ISO | -0.08718 | -0.1997 to 0.02536 | 0.1969 |
| **Δ BEI: Recovery** |  |  |  |
| Tukey's multiple comparisons test | Mean Differences | CI95% adjust | P Value |
| CH vs. KX | -0.04581 | -0.1986 to 0.1070 | 0.9117 |
| CH vs. TBE | 0.04909 | -0.09168 to 0.1899 | 0.8566 |
| CH vs. TP | -0.01463 | -0.1675 to 0.1382 | 0.9987 |
| CH vs. ISO | -0.0918 | -0.2359 to 0.05229 | 0.3784 |
| KX vs. TBE | 0.0949 | -0.05480 to 0.2446 | 0.3835 |
| KX vs. TP | 0.03119 | -0.1299 to 0.1923 | 0.9811 |
| KX vs. ISO | -0.04599 | -0.1988 to 0.1068 | 0.9106 |
| TBE vs. TP | -0.06372 | -0.2134 to 0.08599 | 0.744 |
| TBE vs. ISO | -0.1409 | -0.2817 to -0.0001180 | 0.0497 |
| TP vs. ISO | -0.07718 | -0.2300 to 0.07565 | 0.6065 |

Mean difference = left group − right group. Positive values indicate a higher mean in the first group; negative values indicate a higher mean in the second group.

**Table A6.** Supplementary Statistical Analyses and Results: Blood Gases Biochemical Parameters

| **Δ pH** |  | |  |  |
| --- | --- | --- | --- | --- |
| Tukey's multiple comparisons test | Mean Differences | | CI95% adjust | P Value |
| CH vs. KX | 0.01429 | | -0.07447 to 0.1030 | 0.9899 |
| CH vs. TBE | 0.04571 | | -0.04023 to 0.1317 | 0.547 |
| CH vs. TP | -0.05714 | | -0.1459 to 0.03162 | 0.3586 |
| CH vs. ISO | -0.04554 | | -0.1315 to 0.04040 | 0.5507 |
| KX vs. TBE | 0.03143 | | -0.05451 to 0.1174 | 0.8269 |
| KX vs. TP | -0.07143 | | -0.1602 to 0.01733 | 0.1632 |
| KX vs. ISO | -0.05982 | | -0.1458 to 0.02612 | 0.2838 |
| TBE vs. TP | -0.1029 | | -0.1888 to -0.01692 | 0.0126 |
| TBE vs. ISO | -0.09125 | | -0.1743 to -0.008224 | 0.0255 |
| TP vs. ISO | 0.01161 | | -0.07433 to 0.09755 | 0.9948 |
| **Δ PaCO_2_ (mmHg)** |  | |  |  |
| Tukey's multiple comparisons test | Mean Differences | | CI95% adjust | P Value |
| CH vs. KX | -7.714 | | -20.69 to 5.266 | 0.4382 |
| CH vs. TBE | -8.929 | | -21.50 to 3.640 | 0.2652 |
| CH vs. TP | 0.0 | | -12.98 to 12.98 | >0.9999 |
| CH vs. ISO | -1.679 | | -14.25 to 10.89 | 0.9951 |
| KX vs. TBE | -1.214 | | -13.78 to 11.35 | 0.9986 |
| KX vs. TP | 7.714 | | -5.266 to 20.69 | 0.4382 |
| KX vs. ISO | 6.036 | | -6.532 to 18.60 | 0.6398 |
| TBE vs. TP | 8.929 | | -3.640 to 21.50 | 0.2652 |
| TBE vs. ISO | 7.25 | | -4.892 to 19.39 | 0.4335 |
| TP vs. ISO | -1.679 | | -14.25 to 10.89 | 0.9951 |
| **Δ PaO_2_ (mmHg)** |  | |  |  |
| Tukey's multiple comparisons test | Mean Differences | | CI95% adjust | P Value |
| CH vs. KX | -2.286 | | -20.70 to 16.13 | 0.9963 |
| CH vs. TBE | 16.14 | | -2.268 to 34.55 | 0.1082 |
| CH vs. TP | 17.79 | | -0.04094 to 35.61 | 0.0508 |
| CH vs. ISO | 4.036 | | -13.79 to 21.86 | 0.9646 |
| KX vs. TBE | 18.43 | | 0.01728 to 36.84 | 0.0497 |
| KX vs. TP | 20.07 | | 2.245 to 37.90 | 0.0211 |
| KX vs. ISO | 6.321 | | -11.51 to 24.15 | 0.8421 |
| TBE vs. TP | 1.643 | | -16.18 to 19.47 | 0.9988 |
| TBE vs. ISO | -12.11 | | -29.93 to 5.720 | 0.3068 |
| TP vs. ISO | -13.75 | | -30.97 to 3.472 | 0.169 |
| **Δ HCO_3_ (mmol/L)** |  | |  |  |
| Tukey's multiple comparisons test | Mean Diff, | | 95,00% CI of diff, | Adjusted P Value |
| CH vs. KX | -3.5 | | -8.475 to 1.475 | 0.274 |
| CH vs. TBE | -1.732 | | -6.549 to 3.085 | 0.8353 |
| CH vs. TP | -2.829 | | -7.803 to 2.146 | 0.4822 |
| CH vs. ISO | -3.182 | | -7.999 to 1.635 | 0.3334 |
| KX vs. TBE | 1.768 | | -3.049 to 6.585 | 0.825 |
| KX vs. TP | 0.6714 | | -4.303 to 5.646 | 0.9949 |
| KX vs. ISO | 0.3179 | | -4.499 to 5.135 | 0.9997 |
| TBE vs. TP | -1.096 | | -5.913 to 3.720 | 0.9639 |
| TBE vs. ISO | -1.45 | | -6.103 to 3.203 | 0.8946 |
| TP vs. ISO | -0.3536 | | -5.170 to 4.463 | 0.9995 |
| **Δ Total CO_2_ (mmol/L)** | |  |  |  |
| Tukey's multiple comparisons test | Mean Differences | | CI95% adjust | P Value |
| CH vs. KX | -2.929 | | -7.899 to 2.041 | 0.4467 |
| CH vs. TBE | -1.179 | | -5.991 to 3.634 | 0.9532 |
| CH vs. TP | -2.043 | | -7.013 to 2.927 | 0.7582 |
| CH vs. ISO | -2.441 | | -7.253 to 2.371 | 0.5913 |
| KX vs. TBE | 1.75 | | -3.062 to 6.562 | 0.8297 |
| KX vs. TP | 0.8857 | | -4.084 to 5.856 | 0.9852 |
| KX vs. ISO | 0.4875 | | -4.325 to 5.300 | 0.9983 |
| TBE vs. TP | -0.8643 | | -5.676 to 3.948 | 0.9848 |
| TBE vs. ISO | -1.263 | | -5.911 to 3.386 | 0.9332 |
| TP vs. ISO | -0.3982 | | -5.210 to 4.414 | 0.9992 |
| **Δ O_2_ Saturation (%)** |  | |  |  |
| Tukey's multiple comparisons test | Mean Differences | | CI95% adjust | P Value |
| CH vs. KX | 6.286 | | -1.208 to 13.78 | 0.135 |
| CH vs. TBE | 7.446 | | 0.1908 to 14.70 | 0.042 |
| CH vs. TP | 0.2857 | | -7.208 to 7.779 | >0.9999 |
| CH vs. ISO | 1.196 | | -6.059 to 8.452 | 0.989 |
| KX vs. TBE | 1.161 | | -6.095 to 8.416 | 0.9902 |
| KX vs. TP | -6.0 | | -13.49 to 1.494 | 0.1669 |
| KX vs. ISO | -5.089 | | -12.34 to 2.166 | 0.2767 |
| TBE vs. TP | -7.161 | | -14.42 to 0.09489 | 0.0545 |
| TBE vs. ISO | -6.25 | | -13.26 to 0.7596 | 0.0991 |
| TP vs. ISO | 0.9107 | | -6.345 to 8.166 | 0.9961 |

Mean difference = left group − right group. Positive values indicate a higher mean in the first group; negative values indicate a higher mean in the second group.

**Table A7.** Supplementary Statistical Analyses and Results: Enzymatic Biochemical Parameters

| **Δ Plasmatic BChE Activity (U/L)** | |  |  |
| --- | --- | --- | --- |
| Tukey's multiple comparisons test | Mean Differences | CI95% adjust | P Value |
| CH vs. KX | 7.0 | 1.772 to 12.23 | 0.0042 |
| CH vs. TBE | -0.5 | -5.912 to 4.912 | 0.9988 |
| CH vs. TP | -3.5 | -8.728 to 1.728 | 0.3227 |
| CH vs. ISO | -0.125 | -5.353 to 5.103 | >0.9999 |
| KX vs. TBE | -7.5 | -12.91 to -2.088 | 0.0029 |
| KX vs. TP | -10.5 | -15.73 to -5.272 | <0.0001 |
| KX vs. ISO | -7.125 | -12.35 to -1.897 | 0.0035 |
| TBE vs. TP | -3.0 | -8.412 to 2.412 | 0.5099 |
| TBE vs. ISO | 0.375 | -5.037 to 5.787 | 0.9996 |
| TP vs. ISO | 3.375 | -1.853 to 8.603 | 0.3583 |
| **Δ TnI Concentration (ng/mL)** |  |  |  |
| Tukey's multiple comparisons test | Mean Differences | CI95% adjust | P Value |
| CH vs. KX | 0.6607 | -0.9730 to 2.294 | 0.77 |
| CH vs. TBE | 0.1893 | -1.444 to 1.823 | 0.9972 |
| CH vs. TP | 0.25 | -1.328 to 1.828 | 0.9906 |
| CH vs. ISO | 0.4375 | -1.141 to 2.016 | 0.9289 |
| KX vs. TBE | -0.4714 | -2.159 to 1.216 | 0.927 |
| KX vs. TP | -0.4107 | -2.044 to 1.223 | 0.9491 |
| KX vs. ISO | -0.2232 | -1.857 to 1.411 | 0.9947 |
| TBE vs. TP | 0.06071 | -1.573 to 1.694 | >0.9999 |
| TBE vs. ISO | 0.2482 | -1.386 to 1.882 | 0.992 |
| TP vs. ISO | 0.1875 | -1.391 to 1.766 | 0.9969 |
| **Δ CK-MB Activity (U/L)** |  |  |  |
| Tukey's multiple comparisons test | Mean Differences | CI95% adjust | P Value |
| CH vs. KX | 70.8 | -51.17 to 192.8 | 0.4643 |
| CH vs. TBE | 144.2 | 19.51 to 268.8 | 0.0167 |
| CH vs. TP | 126 | 4.055 to 248.0 | 0.04 |
| CH vs. ISO | 65.43 | -59.24 to 190.1 | 0.5625 |
| KX vs. TBE | 73.38 | -32.88 to 179.6 | 0.2931 |
| KX vs. TP | 55.22 | -47.86 to 158.3 | 0.543 |
| KX vs. ISO | -5.375 | -111.6 to 100.9 | 0.9999 |
| TBE vs. TP | -18.15 | -124.4 to 88.10 | 0.9876 |
| TBE vs. ISO | -78.75 | -188.1 to 30.58 | 0.2545 |
| TP vs. ISO | -60.6 | -166.9 to 45.66 | 0.4819 |
| Δ **LDH Activity** (**U/L**) |  |  |  |
| Tukey's multiple comparisons test | Mean Differences | CI95% adjust | P Value |
| CH vs. KX | 52.15 | -73.19 to 177.5 | 0.7554 |
| CH vs. TBE | 121.0 | -7.980 to 250.0 | 0.0749 |
| CH vs. TP | 83.49 | -41.86 to 208.8 | 0.3304 |
| CH vs. ISO | 24.13 | -104.9 to 153.1 | 0.9829 |
| KX vs. TBE | 68.85 | -56.50 to 194.2 | 0.5225 |
| KX vs. TP | 31.33 | -90.27 to 152.9 | 0.9459 |
| KX vs. ISO | -28.03 | -153.4 to 97.32 | 0.9672 |
| TBE vs. TP | -37.51 | -162.9 to 87.83 | 0.9102 |
| TBE vs. ISO | -96.88 | -225.9 to 32.11 | 0.2201 |
| TP vs. ISO | -59.36 | -184.7 to 65.98 | 0.6577 |

Mean difference = left group − right group. Positive values indicate a higher mean in the first group; negative values indicate a higher mean in the second group.
